# Human-brown bear conflicts in Türkiye are driven by increased human presence around protected areas

**DOI:** 10.1101/2023.08.29.555317

**Authors:** Ercan Sıkdokur, Morteza Naderi, Elif Çeltik, M. Çisel Kemahlı Aytekin, Josip Kusak, İsmail K. Sağlam, Çağan H. Şekercioğlu

## Abstract

Human-wildlife conflict is increasing steadily worldwide and is rapidly becoming an important challenge for the success of conservation programs. Brown bears, which suffer from reduced habitat suitability and quality globally, frequently conflict with humans. These animals need large home ranges to fulfill their habitat requirements. When space and food are restricted, brown bears frequently shift towards human-dominated landscapes that offer reliable food sources. As a country where most of the landscape and habitats are human-dominated, human-brown bear conflict events (HBCs) are frequent in Türkiye. However, there has been no formal analysis of the nature and scope of these conflicts at the country level. Here, using HBC data from 2017 to 2022, we determined the spatial and temporal dynamics of HBC events and generated a risk probability map based on anthropogenic predictors and geographic profiling, to determine the factors driving HBC across Türkiye. HBC events did not show any annual or seasonal trends but varied considerably across biogeographic regions, with most conflicts occurring along the Black Sea coast and Eastern Anatolia. Sixty percent of all conflicts were due to bear foraging behavior in human settlements while twelve percent were the result of human activity in forests, with 57% of all conflict events resulting in direct injury to either humans or bears. We found that distance to villages, distance to protected areas, distance to farmland and human footprint to be the most important factors contributing to conflict risk. Consequently, 21% of the country was found to be under human-bear conflict risk, with 43% of the risks occurring within a 10 km radius from the centers of protected areas. Our analyses indicate that the high occurrence of HBCs is mainly the result of limited natural areas and resources available to brown bears and the increasing human encroachment in and around core bear habitats.

## 1. Introduction

The conservation of large carnivores in human-dominated landscapes is becoming increasingly challenging (Hosseini et al., 2021; Lute et al., 2018). These charismatic animals occupy all trophic levels and have a range of habitat requirements and patterns of space usage that span or require wide landscapes (Habib et al., 2021). Brown bears play a vital role in ecosystems, acting both as ecosystem engineers (Sakiyama et al., 2021) and as megafauna seed dispersers across large areas (García-Rodríguez et al., 2021). Therefore, the preservation of brown bear habitat continuity and quality depends on ecological connectivity across large landscapes, enabling brown bears to utilize a wide range of habitats (Recio et al., 2021).

In the past decade, the expansion of human settlements, and the increase in anthropogenic activities have fragmented numerous ecosystems inhabited by brown bears, resulting in a reduction of brown bear habitats and diminishing the contributions of brown bears to ecosystem services (Morales-González et al. 2020). The expansion of people into the wild animals’ habitats has not only reduced space and resources for these animals but also increased human-wildlife conflict (Dai et al., 2021; Morales-González et al., 2020; Bhandari et al. 2020). Easy access to anthropogenic food sources like open garbage dumps (Cozzi et al., 2016; Plaza and Lambertucci, 2017), orchards (Lamb et al., 2017), and beehives (Naves et al., 2018) aggravate the situation further. Consequently, human-bear conflicts, including attacks on humans (Bombieri et al., 2019), damage to human property (Bautista et al., 2019) and human-caused casualties (i.e., vehicle collisions, illegal hunting etc.) (Gantchoff et al., 2020) are increasing globally (Bombieri et al., 2019). As a result, people and stakeholders can develop negative attitudes toward the conservation of bears, increasing their vulnerability to natural and anthropogenic environmental change (Chapron et al., 2014).

As a 783,562 km^2^ land bridge, Türkiye serves as a major gene flow pathway between Europe, Asia, and Africa. It is almost entirely covered by the Caucasus, Irano-Anatolian, and Mediterranean biodiversity hotspots (Şekercioĝlu et al., 2011a). However, only 13.7% of Türkiye’s area is protected, of which 7.8% is terrestrial (General Directorate of Nature Conservation and Natural Parks of Turkey, 2022), which is lower than the 17% required or recommended by international agreements (Brooks et al., 2004). Furthermore, many of these protected areas are not effectively protected and are threatened by deforestation, construction, mining, other human impacts, and the general undermining of environmental laws (Şekercioĝlu et al., 2011a,b). Although 29% of the country is estimated to be covered by forests (Birben, 2019), various combinations of human activities like logging, farming, construction, and tourism have intensified the ongoing fragmentation and degradation of Türkiye’s forests (Birben, 2019). It is estimated that from 2001 to 2020, Türkiye lost 545,000 ha of its tree cover, equivalent to a 5.4% decrease in total tree cover (Harris et al., 2021). Forestry and forest products play a major role in Türkiye’s economy with export revenues estimated almost 16 billion US dollars end of 2023 (TIM, 2023). The economic importance of Türkiye’s forests makes effective forest management a top priority, but this can also have unintended consequences. For instance, 320,000 km of roads constructed to facilitate forest accessibility (200,000 km of which occur in the forests themselves) has led to both habitat fragmentation and increased public access to natural areas critical for wildlife survival (Bilici, 2021).

With the fragmentation of Türkiye’s forests and the increase in human activity in these areas, Türkiye’s largest carnivore, the Eurasian brown bear (*Ursus arctos arctos*), is facing a wide range of threats, including habitat degradation, poisoning, poaching, and road kills (Ambarli et al., 2016). Additionally, as brown bears are forced to live in increasingly human-dominated landscapes, bear-related incidents within or around human settlements are on the rise, posing a major threat to human-brown bear coexistence. However, we still have little to no information on the pattern and distribution of conflict events across Türkiye and our understanding of the factors that drive or increase conflict risks remains poor. Spatial information on human-brown bear conflicts (HBCs) can help us understand the ecological and anthropogenic factors responsible for these conflicts (Broekhuis et al., 2017) and develop conservation and management strategies to reduce HBCs (Hosseini et al., 2021; Khosravi et al., 2023). Therefore, the aim of this study was to quantify the spatial and temporal trends and determine the social and ecological dynamics of HBCs in Türkiye and outline bear conservation actions that can minimize these conflicts.

## 2. Material and methods

### 2.1. Data collection

To determine the dynamics of the HBC events in Türkiye, we collected and reviewed publicly available data on brown bear-human encounters between 2017-2022. We searched for documented cases of conflict events in publicly available news articles, webpages, official reports and unpublished personal datasets using relevant keywords like ‘bear attacks’, ‘bear mortality’, or a combination of keywords such as ‘crop damage + bear’ in Turkish. All hits were geo-referenced based on the location information. If no location information was available, we mapped results to the nearest location by contacting and confirming with eyewitnesses or the village muhtar (i.e., the local state representative in Turkish villages). If direct eyewitnesses could not be contacted or found, we assigned a random location within 5 km of the event as a georeferenced point (Kimmig et al., 2020).

We classified conflicts into six categories in relation to the nature and cause of the conflict: attack on livestock; attack on beehives; damage to crops; attacks on humans during human activity in forests (gathering resource/food, trekking, camping, logging etc.); traffic accidents involving bears; and illegal killing. Conflicts not falling into these classes were classified as “other”. We employed the Chi-square goodness-of-fit tests to determine significant differences between conflict types and whether the total number of conflicts deviated from a uniform distribution based on conflict type, year, season and across the three major biogeographic regions in Türkiye (Euro-Siberian, Irano-Anatolian, and Mediterranean Davis, 1971; Noroozi et al., 2019).

### 2.2. Environmental layers

We used a combination of anthropogenic variables as environmental predictors, including human population density, housing density, the human footprint index and distance to farmlands, forests, protected areas, roads, villages and water resources (Table S1). Human population and housing density information were extracted from the Socioeconomic Data and Application Center (SEDAC) and OpenStreetMap, respectively. The Human Footprint Index (HFI), which shows the cumulative effect of human pressure on the environment (Venter et al., 2016) was downloaded from the SEDAC website. Forest and farmland layers were extracted from the Corine Land Use Land Cover 2018 vector dataset. Motorways, primary, secondary and trunk roads and their linked attributes were selected and extracted from the Open Street Map. For the rivers, we used the HydroRIVERS database and extracted 1^st^ to 4^th^ order river attributes (Lehner and Grill, 2013). The protected areas layers were downloaded from the GeoData application of the General Directorate of Nature Conservation and National Parks of Türkiye (MOAF, 2019). The following categories of protected areas were extracted and used in our modeling approach: national parks; nature conservation areas; special environment protection areas; wildlife enhancement areas (equivalent to wildlife refuges or reserves); locally important wetlands; nationally important wetlands; RAMSAR wetlands; and preservation (protected) forests. Finally, we generated distance layers for all our variables using the Euclidean Distance tool in ArcGIS (ESRI Inc, USA).

We used the Pearson correlation coefficient and the variance inflation factor to check for multi-collinearity between the nine predictors (Dormann et al., 2013). We did not find any significant multi-collinearity between predictors as all pairwise correlations were below |r|<0.75 and the variance inflation factor for all variables was below 5 (Elith et al., 2006). Therefore, all nine variables were used as input in our modeling analyses (Table. S1).

### 2.3. Human - Brown bear conflict risk modeling

Human-brown bear conflict risk modeling was based on 211 geo-referenced conflict records (Fig. 1). To avoid spatial autocorrelation, we calculated the relatedness of environmental predictors within the occurrence records via a Mantel test using the *ecospat* package in R (Boavida et al., 2016; Broennimann et al., 2018). If there was more than one occurrence record at a distance shorter than the minimum significant autocorrelated distance (14.44 km), only one record was used in the analysis (Fig. S1). Ultimately, we retained 140 occurrence points that were used to model human-brown bear conflicts.

**Fig. 1.**
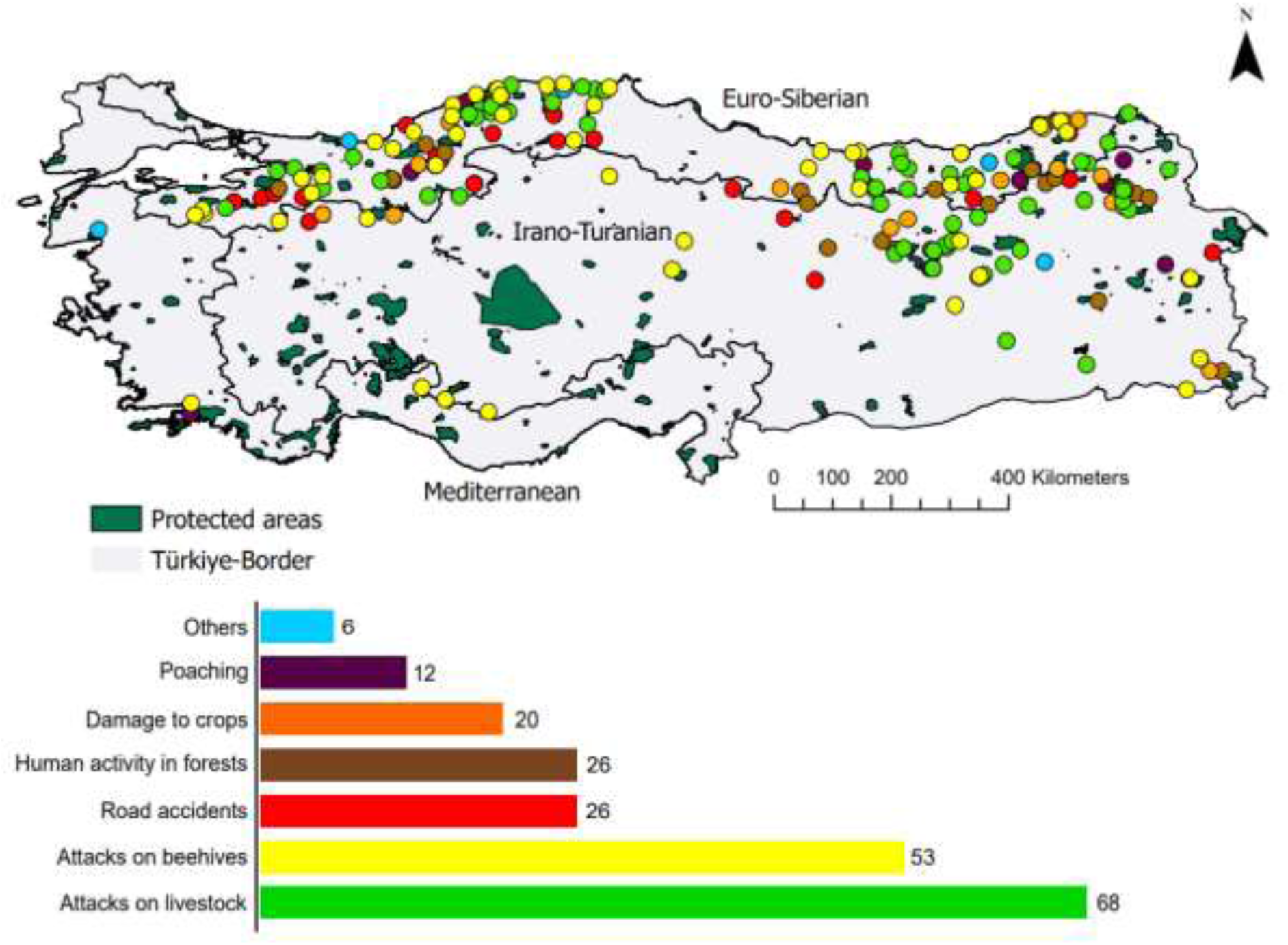
Distribution and number of recorded conflict events and types across Türkiye between 2017-2022.

We used Ecological Niche Modeling (ENM) to determine the distribution of human-brown bear conflict risk probabilities across Türkiye using the *biomod2* package in R (Thuiller, 2014). Specifically, we employed eleven different ENM models based on the following algorithms: 1) Generalized Linear Model (GLM) (P. McCullagh & Nelder, 1989), 2) Generalized Additive Model (GAM) (Hastie and Tibshirani, 2017), 3) Maximum Entropy (MAXENT) (Phillips et al., 2004), 4) Random Forest (RF) (Breiman, 2001), 5) Artificial Neural Network (ANN) (Lek and Guégan, 1999), 6) Multivariate Adaptive Regression Spline (MARS) (Leathwick et al., 2005), 7) Generalized Boosted Model (GBM) (De’ath, 2007), 8) Classification Tree Analyses (CTA) (Vayssières et al., 2000), 9) Maxnet (Phillips et al., 2017), 10) Flexible Discriminant Analysis (FDA) (Hastie et al., 1994), and 11) Surface Response Envelope (SRE) (Busby, 1991). Modeling was performed by generating 1,000 random pseudo-absence points and splitting the collected data into training (80%) and testing (20%) datasets (Pant et al., 2021). To avoid sampling bias, pseudo-absence points were generated three times for each model and cross-validated per model using bootstrapping (Carvalho et al., 2021). We evaluated and compared the performance and accuracy of each model using AUC (area under the receiver operating curve) and TSS (true skill statistic) (Araújo and New, 2007). True skill statistics value changes between −1 and +1, with positive values indicating a perfect fit, while AUC values range from 0.5 to 1, with higher values indicating improved accuracy and predictive power (Fielding and Bell, 1997).

To determine risk probabilities across Türkiye and to reduce the uncertainty among models, we constructed a final ensemble conflict risk model by weight averaging the eleven different models as implemented in the *biomod2* package (Thuiller, 2014). By combining predictions from several different modeling approaches, ensemble ecological niche models have been shown to produce more robust results when compared to single niche models (Araújo and New, 2007; Beeman et al., 2021). We averaged predictions across three replicated runs for each model and selected model runs with a TSS > 0.5 for generating the ensemble model. We classified the predicted risk probabilities generated by the ensemble model into 5 classes: no-risk areas (< 0.2), low-risk areas (0.2-0.4), moderate-risk areas (0.4-0.6), high-risk areas (0.6-0.8) and extreme risk areas (> 0.8). To better understand conflict risk probabilities and human activity around protected areas we created two separate buffered layers around each protected area: a layer spanning an area of 1) 5km radius and 2) a 10 km radius from the center of each protected area. We then determined conflict risk probabilities, housing density and the HFI index within these layers. Finally, we presented the contribution of each variable to our ensemble model and used response curves to visualize the relationship of each variable to the predicted risk assessment.

## 3. Results

### 3.1. Spatio-temporal patterns of human-brown bear conflicts

We determined 211 human-brown bear conflicts in Türkiye between 2017 and 2022 (Fig. 1). Across the six years, the annual number of HBC events averaged 35.16 ± 15.51 (range: 25-65). Except for a significant increase in 2021 (n=65, *X^2^ =* 35.14; df=1, *P* = 3.07e-09), the number of yearly HBC events did not deviate significantly from the mean (Fig 2, Table S2). As expected, the number of conflict events was low in winter, corresponding to the hibernation period of brown bears, and increased significantly in spring but did not show any significant differences among the spring, summer and fall seasons (*X^2^ =* 2.3077, df = 2, *P* = 0.3154) (Fig. 2, Table S3). We conclude that overall HBC events across Türkiye did not show any significant annual or seasonal trends.

**Fig. 2.**
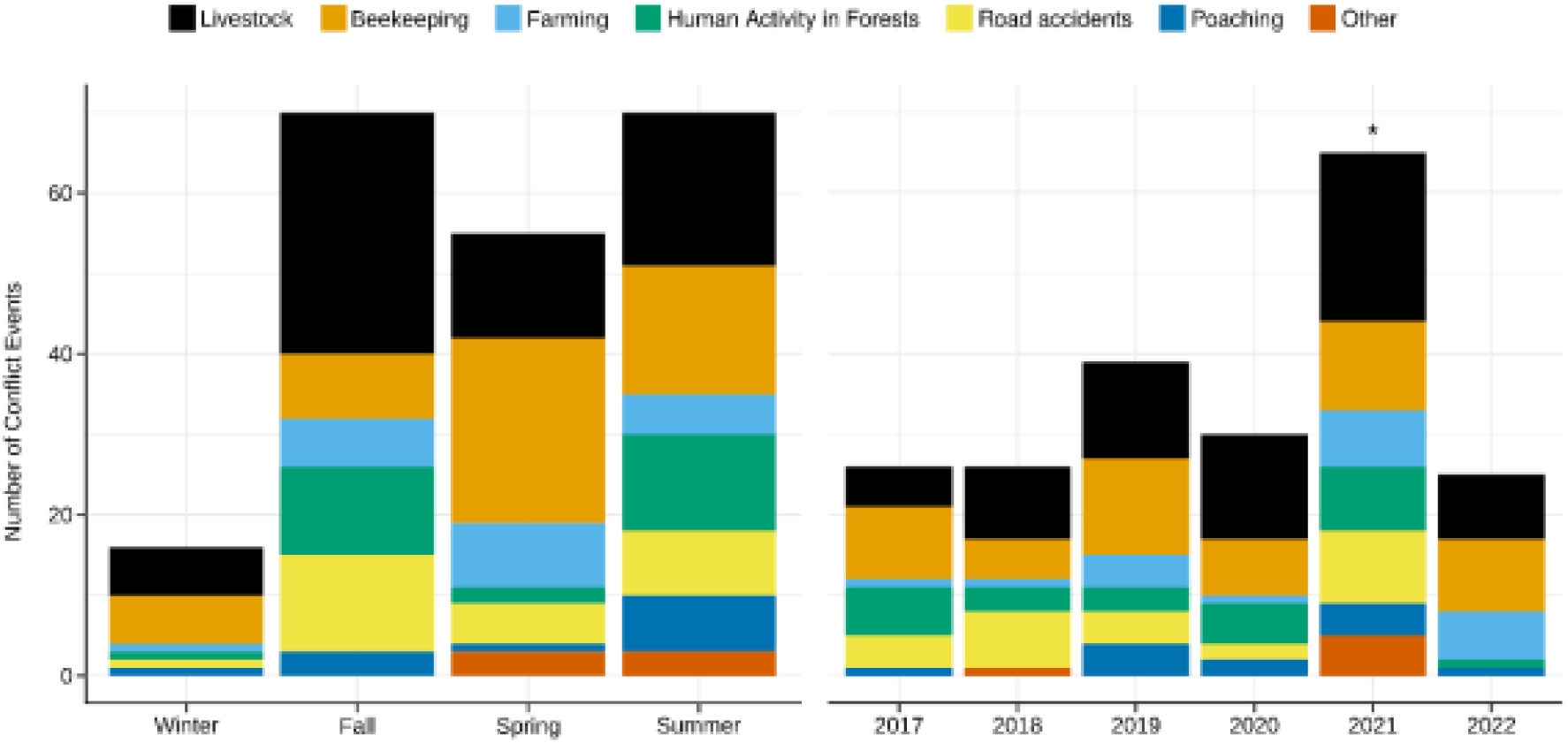
Number of conflict events across seasons and years from 2017-2022 in Türkiye. *Denotes statistically significant deviation from the mean based on a Chi-square test of goodness of fit.

Differences among the frequencies of different conflict types (excluding “others”) throughout Türkiye were highly significant (*X^2^* = 99.687, df = 5, *P* < 2.2e-16) (Table 1). Attacks on livestock (n=68) represented the highest number of conflict events followed by damage to beehives (n=53), damage to crops (n=20), human activity in forest (n=26), traffic accidents (n=26) and illegal killing (n=12) (Table 1). The total number of conflicts showed considerable variation across biogeographic regions (Table 1) but did not correlate with the size of the area (*r* = 0.07; *P* = 0.95). Despite having the smallest area, the highest number of conflicts were observed along the Caucasus and West Black Sea habitats of the Euro-Siberian region (n=130, 146,694 km^2^, 0.080 conflict events per 100 km^2^), and was significantly higher than the number of events recorded in the Irano-Turanian region (n=77, 480,937 km^2^, 0.017 conflict events per 100 km^2^) (X2 = 13.570; df=1; P = 2.30e-04). In contrast, very few conflict events were observed in the Mediterranean region (n=4, 153,073 km^2^, 0.003 conflict events per 100 km^2^) (Table 1). The frequency of occurrence of conflict types also differed between the Euro-Siberian and Irano-Turanian regions as attacks on livestock, attacks on beehives and road accidents were significantly higher in the Euro-Siberian region (Table 1).

**Table 1.**
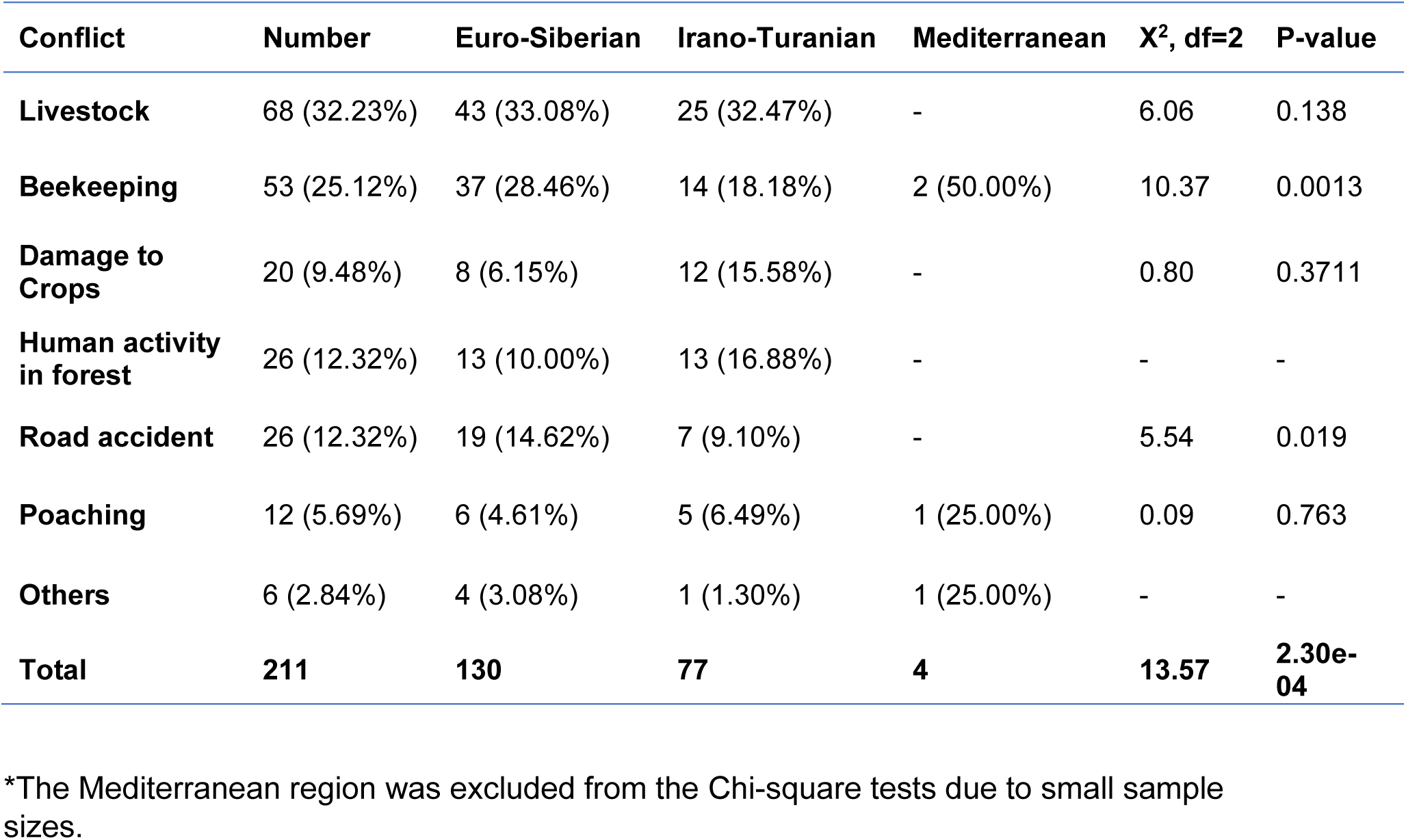
The number and frequency of human-brown bear conflict events within Türkiye and the three biogeographic regions, including the results of Chi-square tests* of the null hypothesis that the percentage contribution of each conflict type does not differ between the Euro-Siberian and Irano-Turanian regions.

| Conflict | Number | Euro-Siberian | Irano-Turanian | Mediterranean | X<sup>2</sup>, df=2 | P-value |
| --- | --- | --- | --- | --- | --- | --- |
| **Livestock** | 68 (32.23%) | 43 (33.08%) | 25 (32.47%) | - | 6.06 | 0.138 |
| **Beekeeping** | 53 (25.12%) | 37 (28.46%) | 14 (18.18%) | 2 (50.00%) | 10.37 | 0.0013 |
| **Damage to Crops** | 20 (9.48%) | 8 (6.15%) | 12 (15.58%) | - | 0.80 | 0.3711 |
| **Human activity in forest** | 26 (12.32%) | 13 (10.00%) | 13 (16.88%) | - | - | - |
| **Road accident** | 26 (12.32%) | 19 (14.62%) | 7 (9.10%) | - | 5.54 | 0.019 |
| **Poaching** | 12 (5.69%) | 6 (4.61%) | 5 (6.49%) | 1 (25.00%) | 0.09 | 0.763 |
| **Others** | 6 (2.84%) | 4 (3.08%) | 1 (1.30%) | 1 (25.00%) | - | - |
| **Total** | **211** | **130** | **77** | **4** | **13.57** | **2.30e-04** |

\*The Mediterranean region was excluded from the Chi-square tests due to small sample sizes.

Among the 211 recorded conflict events, 79 (37.4%) included direct attacks on humans, which left 69 people injured and 12 people dead. The majority of the conflict events resulting in attacks on humans were because of human activities in forests (n=59, 74.7%) including animal husbandry (n=34), picking mushrooms (n=9), hiking (n=8), picking berries (n=4), beekeeping (n=2), and logging (n=2) but attacks on humans in farmlands or crop areas were also quite common (n=14, 17.7%). Conversely, 43 conflict events (20.4%) resulted in brown bear casualties, which left 14 bears injured and 33 dead. The main conflict events causing brown bear casualties were traffic accidents (n=24) and illegal killing (n=11).

### 3.2. Conflict modeling and Anthropogenic factors affecting human-brown bear conflicts

Among the ten models we evaluated, all except SRE (AUC = 0.64) had good to excellent performance based on AUC scores (AUC = 0.79 - 1.00) whereas performance based on TSS values was much more variable and ranged from poor (TSS < 0.4) to good (0.6 < TSS < 0.8) to excellent (TSS > 0.8) (Fig. S4). The best-supported models based on both AUC and TSS were RF (AUC = 1; TSS = 1), GBM (AUC = 0.95; TSS = 0.79) and Maxent (AUC = 0.92; TSS = 0.65) (Fig. S4). On the other hand, the weighted average of AUC and TSS scores across all eleven models were 0.87 and 0.62, respectively, indicating that the ensemble model had good discriminatory power overall (Fig. S4). Based on the ensemble model, the top four factors contributing to conflict risk were distance to villages (37%), distance to protected areas (18%), distance to farmland (16%) and human footprint (13%) (Fig. 3). Response curves of the top four predictors showed that conflict risk decreased with increasing distance to villages and protected areas but showed a bell curve relationship with distance to farmland and human footprint (Fig. 4).

**Fig. 3.**
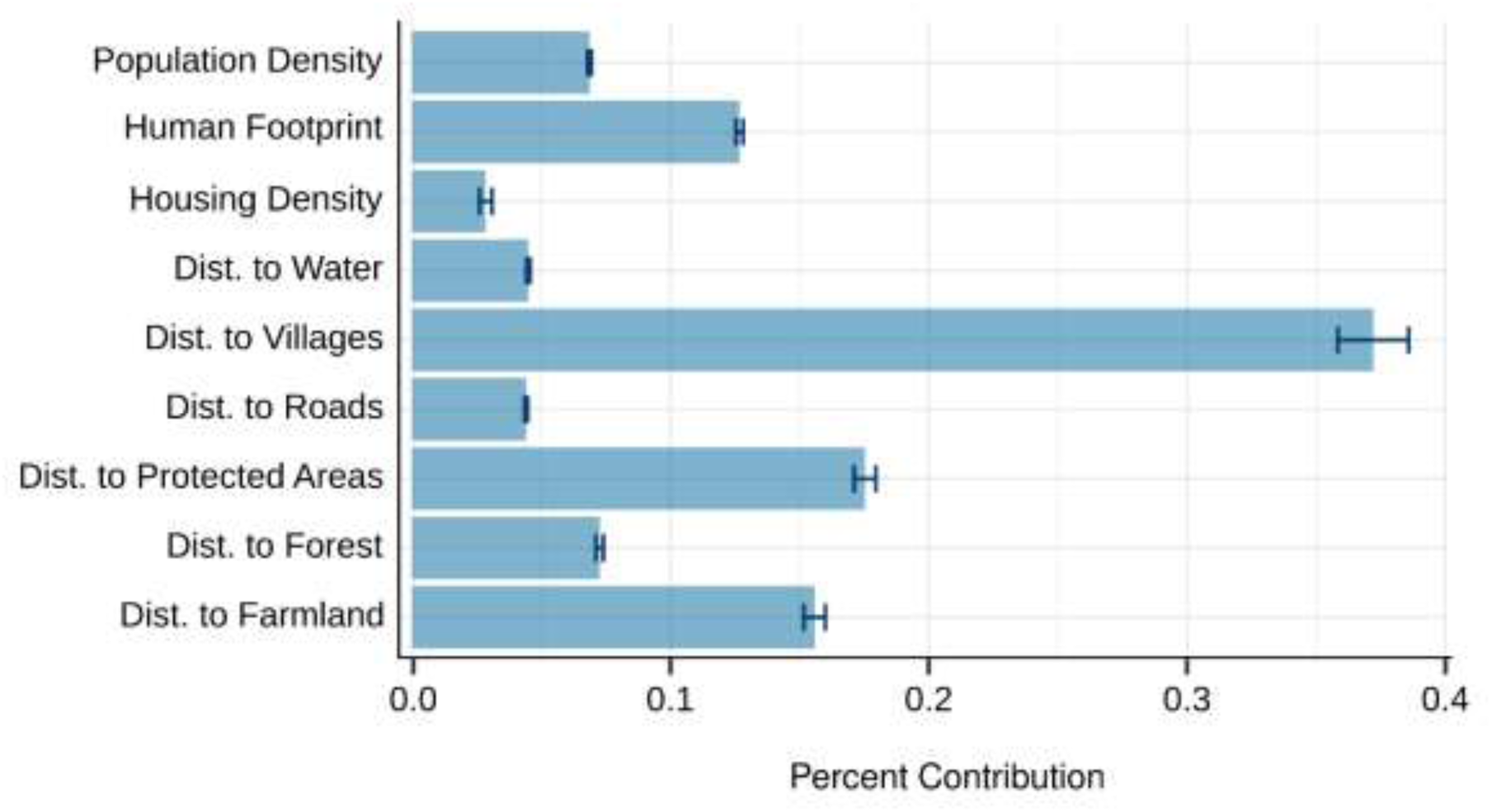
Percent contribution of anthropogenic variables to the final ensemble model for predicting human-brown bear conflicts across Türkiye.

**Fig. 4.**
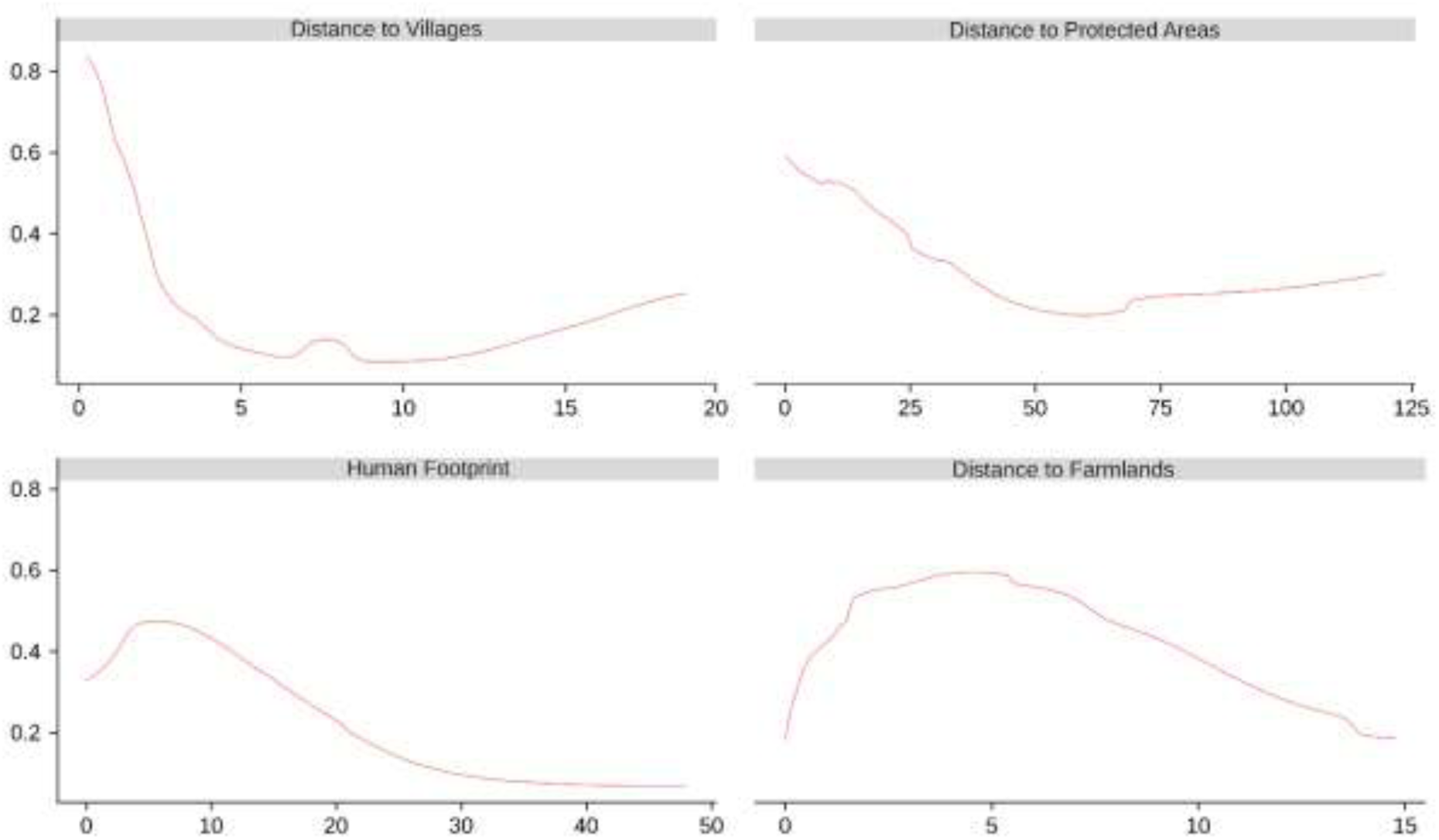
Response curves of the top 4 anthropogenic variables in the ensemble model for predicting human-brown bear conflicts across Türkiye.

**Fig. 5.**
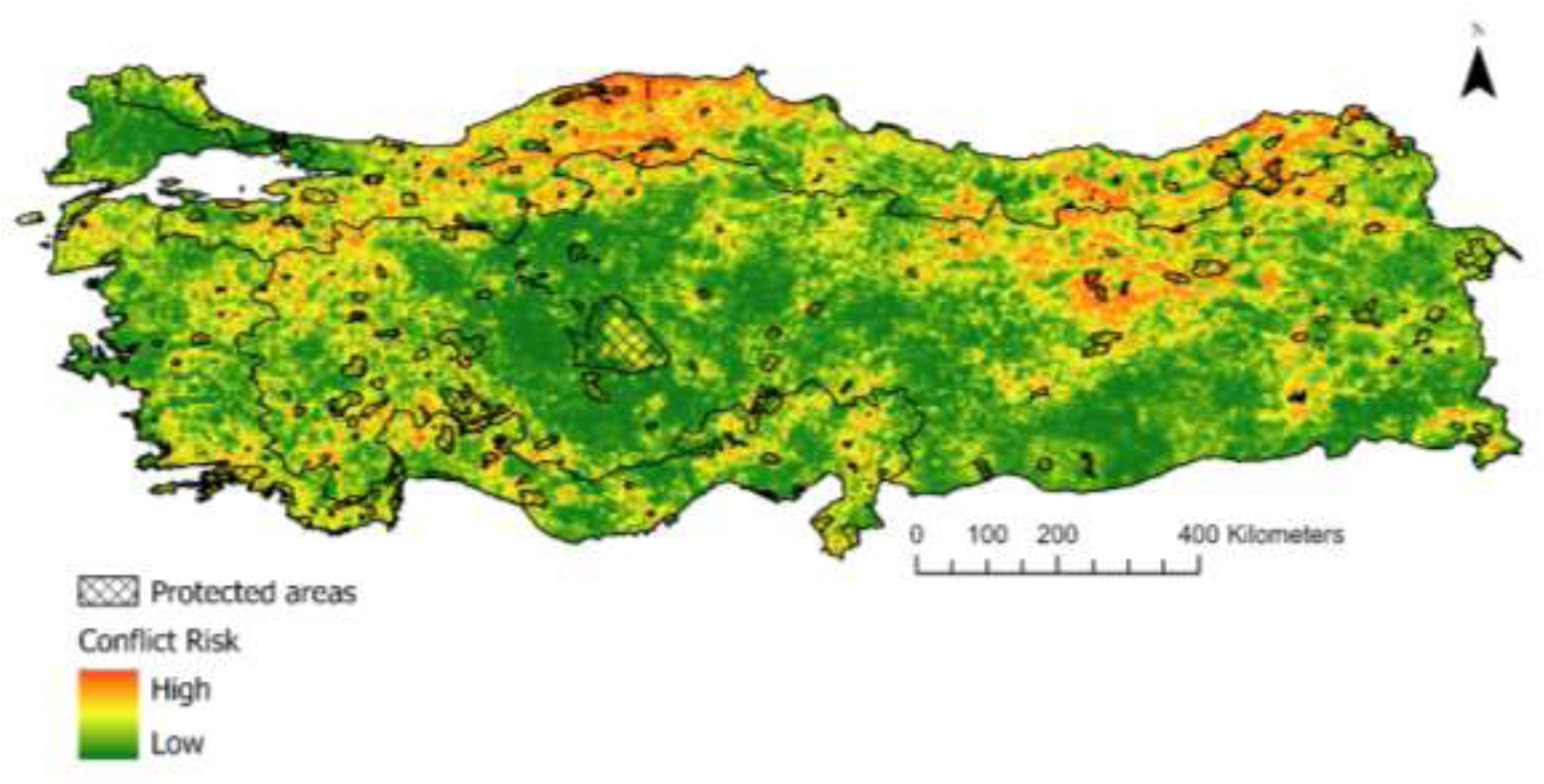
Ensemble model for risk probability of human-brown bear conflict across Türkiye.

A total of 164,311 km^2^ of area (21% of the country) was determined to be under human-brown bear conflict risk. High-risk areas are mostly concentrated along the Black Sea coast and in Eastern Türkiye (Fig. 6) and 7.5% of conflict-risk areas are inside protected areas (Fig. S6). 24.5% (191,780 km^2^) and 14.2% (111,188 km^2^) of the country are classified as low to moderate risk while 8.47% (66,327 km^2^) and 3.3% (25,729 km^2^) are classified as high to extreme risk (Fig. 6). The two buffered layers circling a 5 and 10 km radius from the center of protected areas shows that 24% of all conflict risk areas are within a 5 km radius around protected areas and this increases to 43% within a 10 km radius (Fig. S6). Buffered layers also revealed high human activity (Human Footprint Index > 10) and a high number of human settlements (>6,000 settlements) around protected areas (Table S4).

**Fig. 6.**
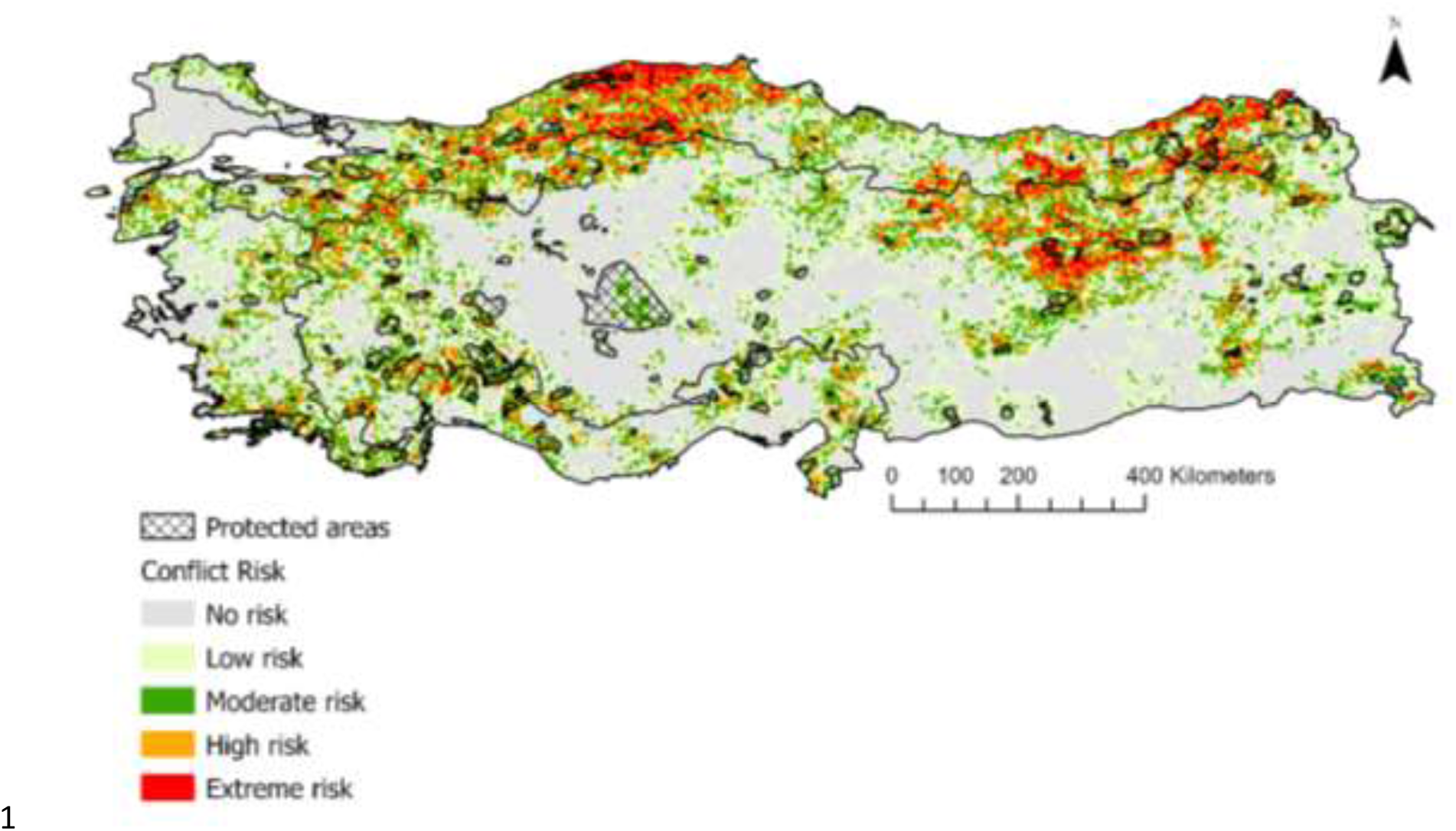
The scaled conflict risk probability across the country.

## 4. Discussion

To our knowledge, this is the first comprehensive study to document and assess nationwide patterns of human-brown bear conflicts (HBCs) in Türkiye. The brown bear is one of Türkiye’s most abundant and widely distributed large mammals, and human-brown bear conflicts are a common occurrence across the country (Can and Togan, 2004). Several factors have been proposed for explaining the frequent occurrence of HBCs in Türkiye, including habitat fragmentation and patchiness, increased development and construction in forest habitats and higher bear numbers especially along the Black Sea region of Türkiye (Ambarlı et al., 2018). Here, using HBC data from 2017 to 2022 we determined the spatial and temporal dynamics of HBC events and generated a probability risk map based on anthropogenic predictors and geographic profiling to determine factors driving human-brown bear conflicts across Türkiye.

Human-wildlife conflicts are common whenever humans and animals share the same space (Bombieri et al., 2023; Nayeri et al., 2022; Wahab et al. 2021). Our data is also reflective of this observation, as high conflict risk areas and most of the conflicts (98%) occurred along the Black Sea coast and Eastern Anatolia (Fig. 6), where human settlement and activity have historically been associated with rural and forested areas (Koc et al., 2015). Although no formal study has censused bear numbers in Türkiye, the Black Sea and Eastern Anatolia regions are known to support relatively large bear populations, with the total number of individuals across both regions estimated to be around 2000-3000 (Ambarli et al., 2016). In contrast, the Mediterranean region, estimated to support around 200 to 300 bears, had little to no record of human-bear conflicts (n=4) and was generally classified as a low-conflict risk area (Fig 6).

While these results show a link between high bear numbers and the frequency of conflicts, prior studies show that reducing population sizes (i.e., harvesting) has little impact on the levels of HBCs, especially in regions where the availability of natural foods and foraging space is restricted (Acharya et al., 2017; Bautista et al., 2023). When space and food are restricted in natural habitats, wide-ranging mammals and especially bears, can shift towards human-dominated landscapes which offer predictable and reliable sources of food with reduced foraging cost (Bautista et al., 2023; Kemahlı Aytekin, 2022). Protected areas in Türkiye represent little coverage of natural bear habitats, with only 6.3% and 4.6% of all forested areas within the Black Sea region and East Anatolia being protected based on the Corine database, and some of these forests are legally logged. The immediate vicinity of protected areas is also heavily populated in Türkiye, with 26% of all human settlements located within a 10 km radius of protected areas (Table S4). Therefore, it is not surprising that brown bears, which have typical home ranges of around 14-83 km^2^ in Türkiye (Ambarli et al., 2016), interact heavily with human settlements around their core habitats for space and natural resources. Our models show that the risk of conflicts is relatively low within protected areas (7.8%), but this number increases dramatically (43%) within a 10 km radius around protected areas. Indeed, the most important factor associated with HBCs in Türkiye was the distance to villages (37%), with distance to farmland (16%) also having a strong contribution (Fig. 3). These results indicate the heavy reliance on Turkish brown bears on human resources, as 67% of all conflicts in our data set were related to foraging activities in human settlements (attack on livestock 32.23%, attack on beehives 25.12%, and damage to crops 9.48%). Some conflicts resulting from brown bear foraging activities, such as damages to orchards, are also a common occurrence in Turkish villages, although they are far less featured in media coverage. Therefore, if anything, our data set is an underestimate of human-brown bear conflicts caused by bear foraging activity within human settlements.

Apart from distance to villages and farmland, the other most important factors explaining patterns of HBCs in Türkiye were distance to protected areas (18%) and the human footprint index (13%) (Fig. 3). This can be reflective of the high amount of human activity in and around core protected areas and forests, which was responsible for 12.32% of all conflicts in Türkiye (Table 1). Although, human-caused conflicts resulted mainly from direct human activities in forested areas like logging, poaching, and recreation (i.e. trekking, picking berries, gathering mushrooms, camping, etc.), accidental road kills also amounted to an important percentage (Table 1). Overall, the human footprint is highly elevated within and around protected areas in Türkiye (Table S4), with an average HFI of 9.81 ± 5.80 (Table S4). Given a typical HFI score of ≥6 for human settlements (Venter et al., 2016), this number is way too high for areas that should be heavily regulated for human activity. High amounts of human activity within and around core protected areas can also explain the relatively high attack rates of brown bears on humans, as 37.44% of all conflicts in Türkiye resulted in some form of altercation. The annual attack rate on humans in Türkiye was calculated to be 13.2 attacks per year, which is higher than continent-wide attack rates in North America (11.4 attacks/year) and Europe (10 attacks/year, excluding Romania) (Bombieri et al., 2019). Apart from human casualties, bear casualties were also high, with 20.37% of all encounters ending with harm to bears. The majority of these were the result of road accidents (n=24). Roads are major ecological traps in many parts of the world and can lead to increased human-wildlife conflict by restricting wildlife movement and concentrating human activity in otherwise wild or natural areas (Lamb et al., 2017). The situation in Türkiye is much the same, as most road accidents occurred in the Black Sea region, where roads increasingly bisect forests and where road construction in forested areas is steadily increasing. To a lesser degree, poaching is also of some concern in Türkiye, as we documented 11 cases of brown bear mortality due to illegal hunting. This is likely to be an underestimate because hunting brown bears is illegal in Türkiye.

## 5. Conclusion

Large mammals like brown bears, which are increasingly driven to survive in human-dominated landscapes (Morales-González et al., 2020) often cannot persist inside small conservation areas and critically depend on external habitats for food and other resources. This tension creates opportunities for increased human-wildlife conflict, especially if human activity within and around protected areas is rampant and unregulated. As our study shows, this seems to be the main reason behind elevated HBC numbers in Türkiye. The core habitats and movement corridors of brown bears increasingly bisect human settlements in Türkiye, and human activity is largely unchecked around these areas. Moving forward, identifying core brown bear habitats and movement corridors that can meet the requirements of this species and establishing newly expanded borders for protected areas should be the main priority of conservation efforts if we want to minimize human-wildlife conflicts. Furthermore, our study shows that human activity within protected areas, such as logging, collecting, livestock grazing, and recreational activities, needs to be more strictly regulated, especially if we want to reduce unwanted outcomes such as human or brown bear mortality. Our results suggest that the high occurrence of HBCs in Türkiye is mainly the result of limited protected area coverage, natural habitats and resources available to brown bears and increasing human encroachment in and around core wildlife habitats.

## Credit authorship contribution statement

**ES, MN, CS, IKS** conceptualized the study**. ES, MCKA, EC** collected and organized the data**. ES, MN** ran all formal analyses with input from **IKS. ES, MN, IKS** wrote the manuscript with input from **JK** and **CS.**

## Declaration of competing interest

The authors declare that there are no conflicts of interest.

## Acknowledgement

The study was supported by The Scientific and Technological Research Council of Turkey (TUBITAK), project number 219Z066.

## Supplementary Material

**Figure S1:**
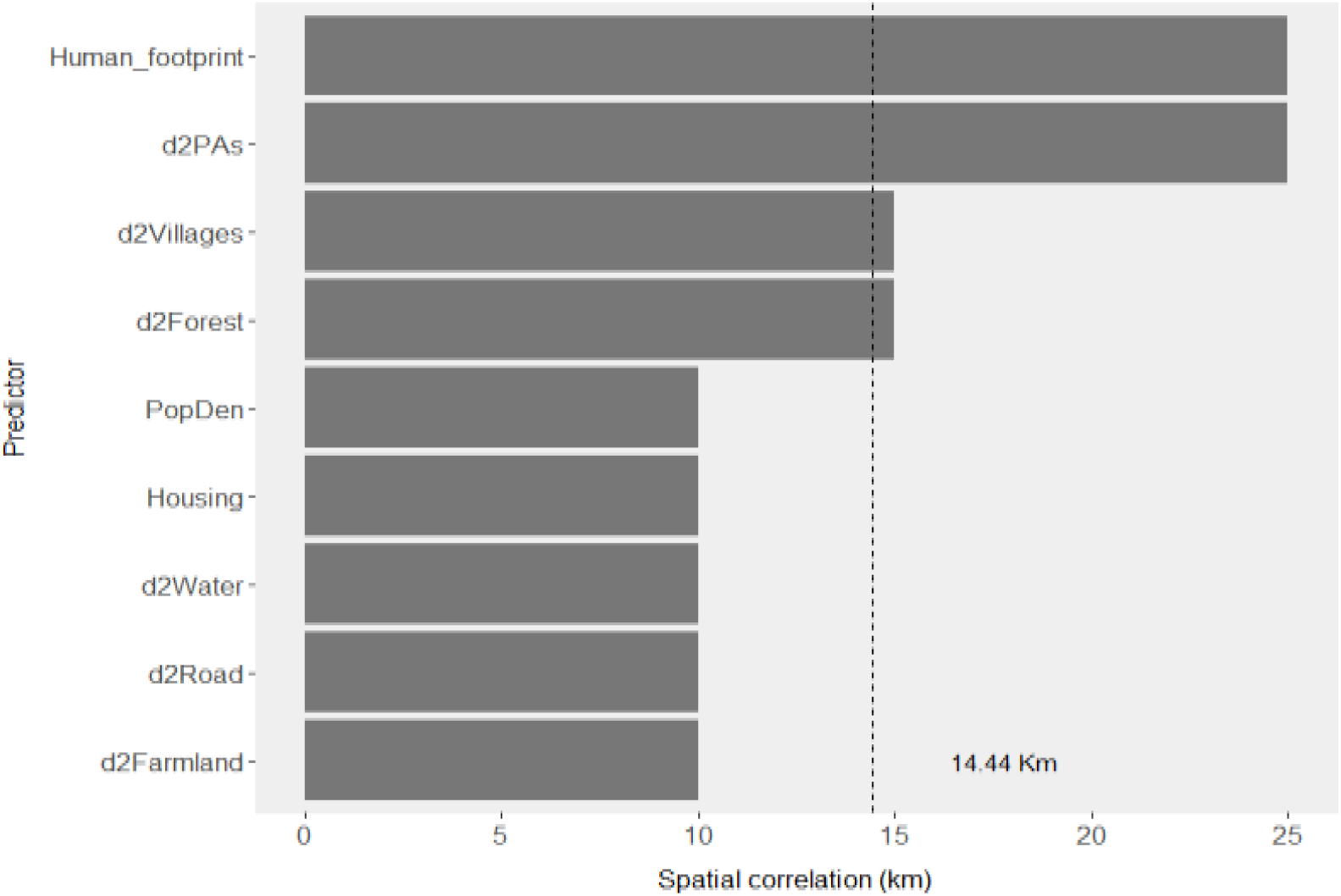
Minimum and average (dashed line) non-significant autocorrelated distances of variable predictors. d2Farmland: Distance to farmlands; d2Forest: Distance to forests; d2Road: Distance to roads; d2PAs: Distance to protected areas; d2Villages: Distance to villages; d2Water: Distance to waters; Human_footprint: Human Footprint Index; PopDen: Population Density; Housing: Housing Density.

**Figure S2:**
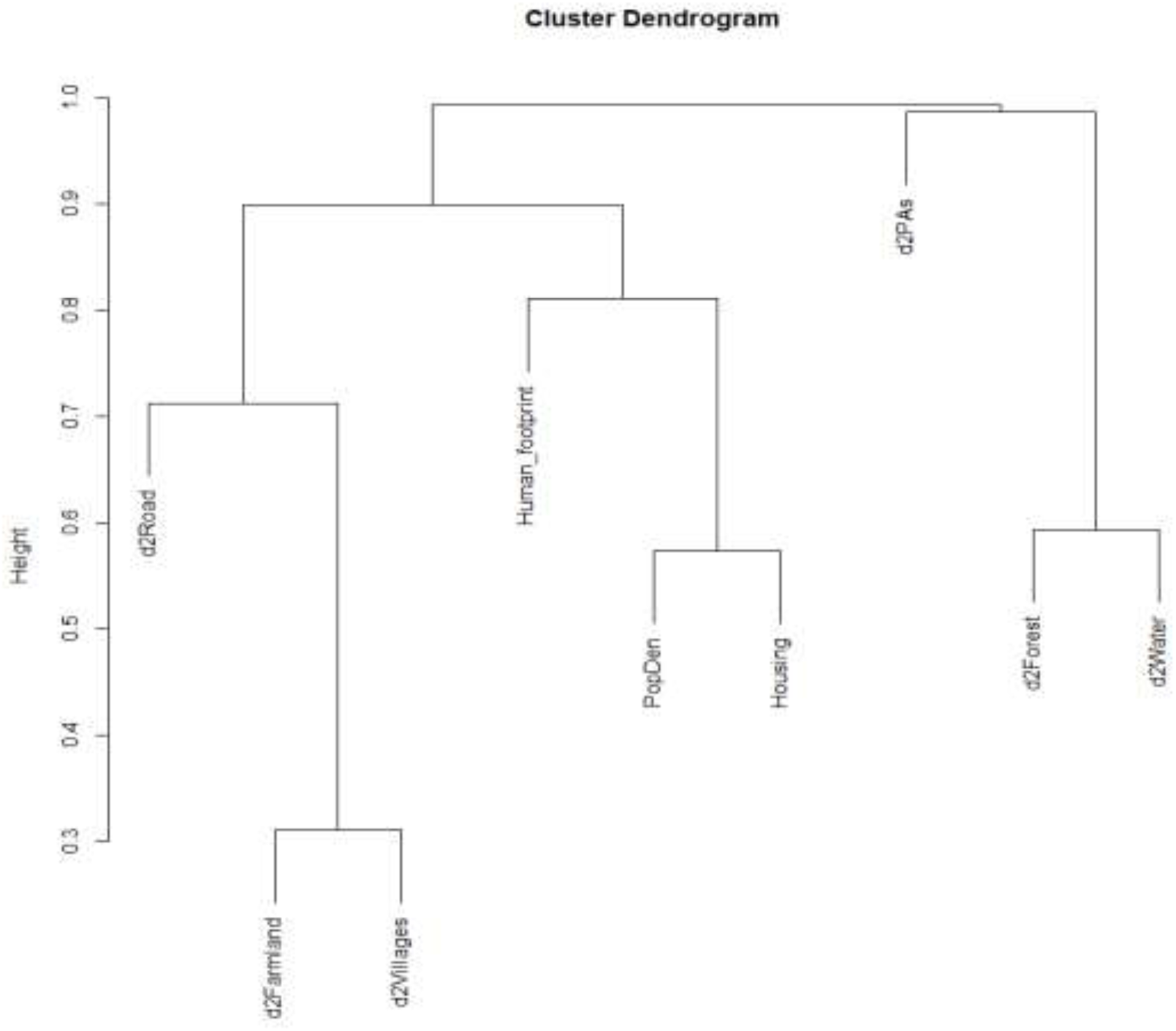
Cluster dendogram of the environmental variables. d2Farmland: Distance to farmlands; d2Forest: Distance to forests; d2PAs: Distance to protected areas; d2Road: Distance to roads; d2Villages: Distance to villages; d2Water: Distance to waters; Human_footprint: Human Footprint Index; PopDen: Population Density; Housing: Housing Density.

**Figure S3:**
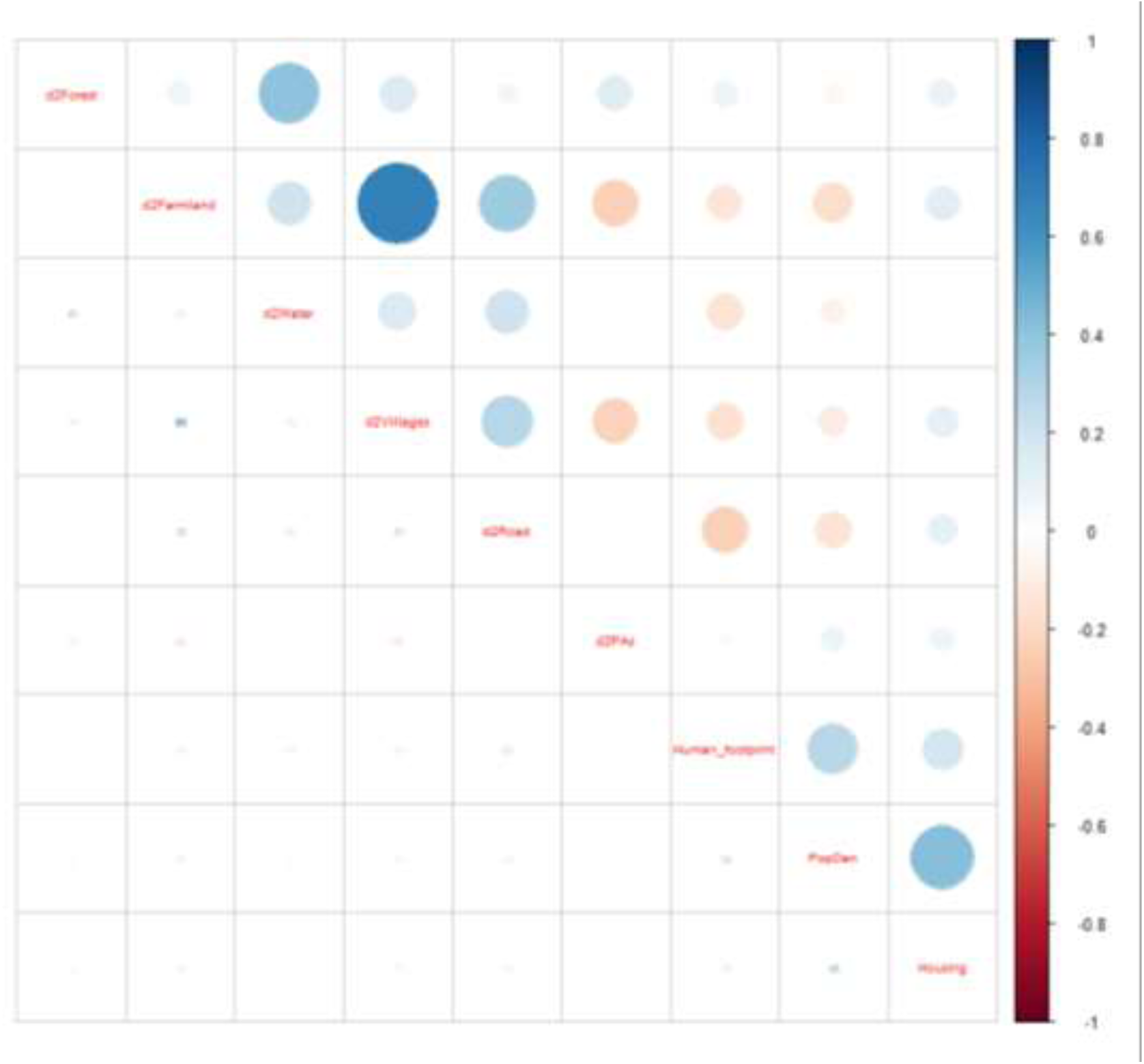
Correlation matrix of the environmental variables. d2Farmland: Distance to farmlands; d2Forest: Distance to forests; d2PAs: Distance to protected areas; d2Road: Distance to roads; d2Villages: Distance to villages; d2Water: Distance to waters; Human_footprint: Human Footprint Index; PopDen: Population Density; Housing: Housing Density.

**Figure S4:**
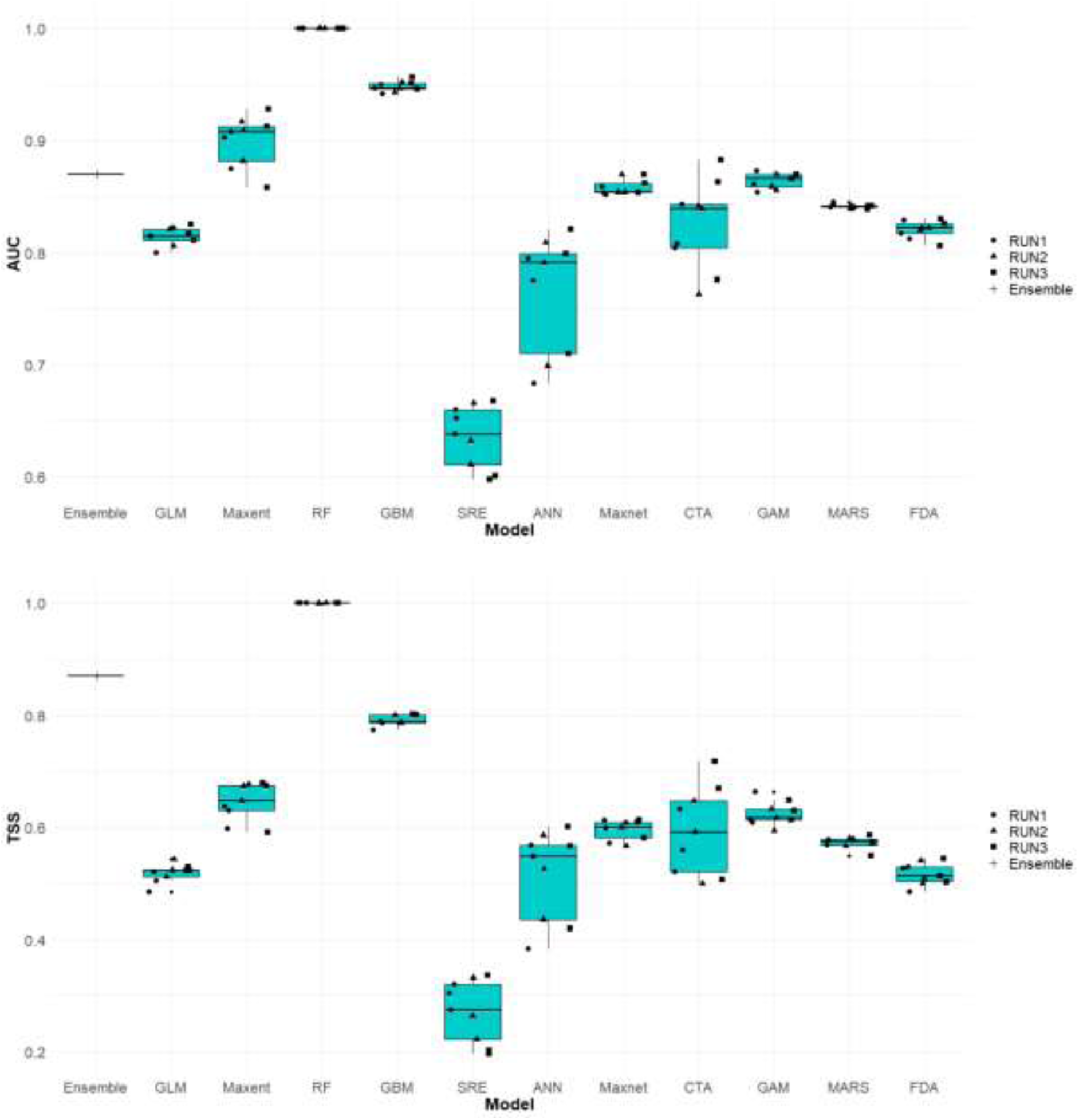
Receiver operating characteristic (ROC) area under the curve (AUC) (above), and True-skill statistics (TSS) (below) values as predictive performance of different modeling techniques.

**Figure S5:**
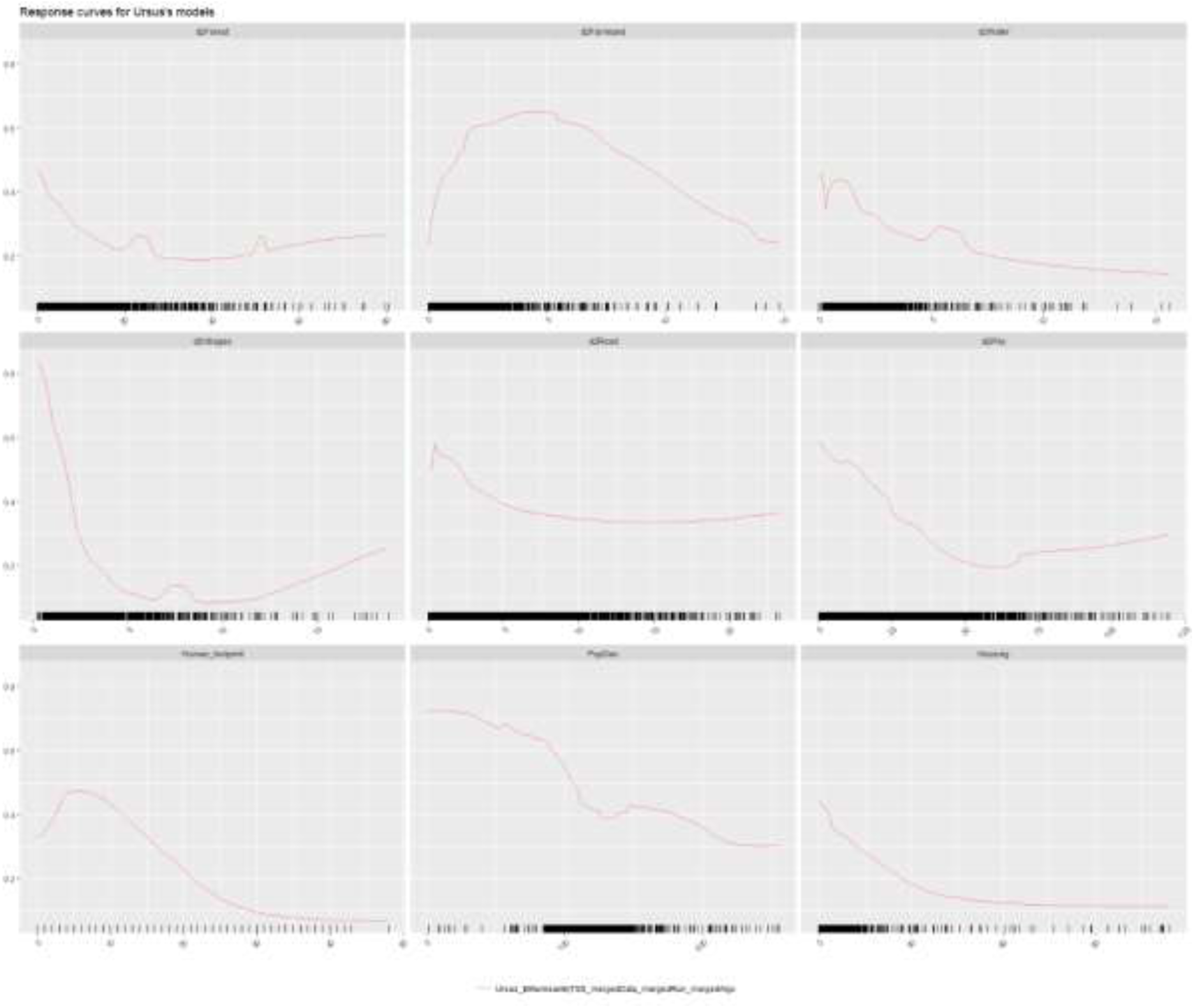
Response curves of the environmental variables used to risk modeling of human-brown bear conflict across the country. d2Farmland: Distance to farmlands; d2Forest: Distance to forests; d2PAs: Distance to protected areas; d2Road: Distance to roads; d2Villages: Distance to villages; d2Water: Distance to waters; Human_footprint: Human Footprint Index; PopDen: Population Density; Housing: Housing Density.

**Figure S6:**
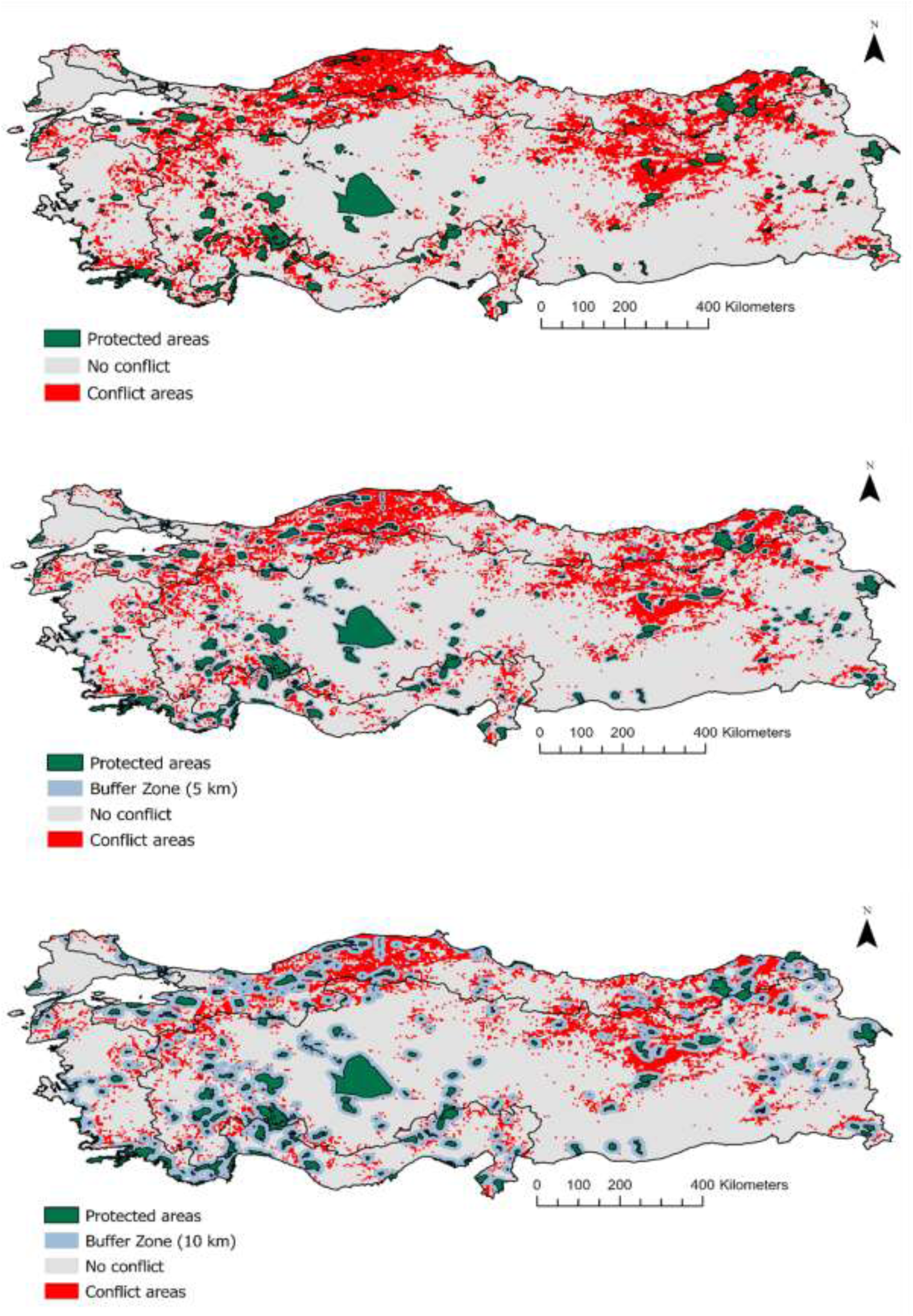
Binary map of conflict risk across Türkiye based on the ensemble model for 0 (top), 5 (mid) and 10 km buffer zone (bottom) around protected areas.

**Table S1:**
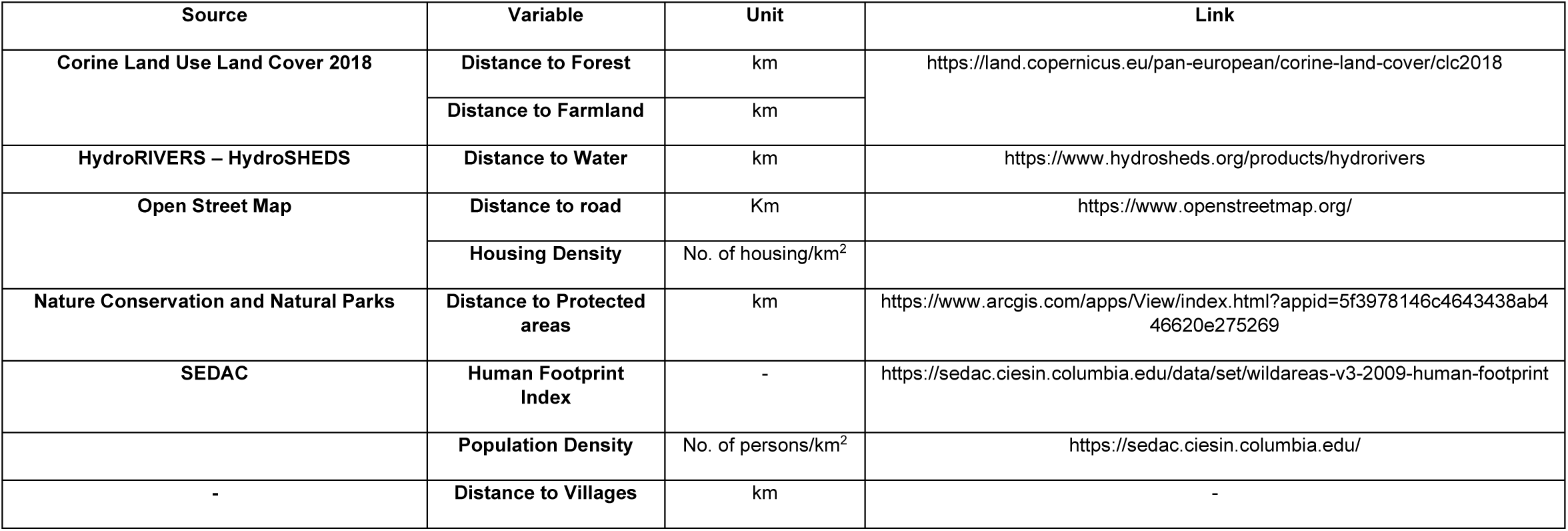
Environmental variables used for risk modeling of human-brown bear conflict across Türkiye. All variables were selected to run final model.

**Table S2:**
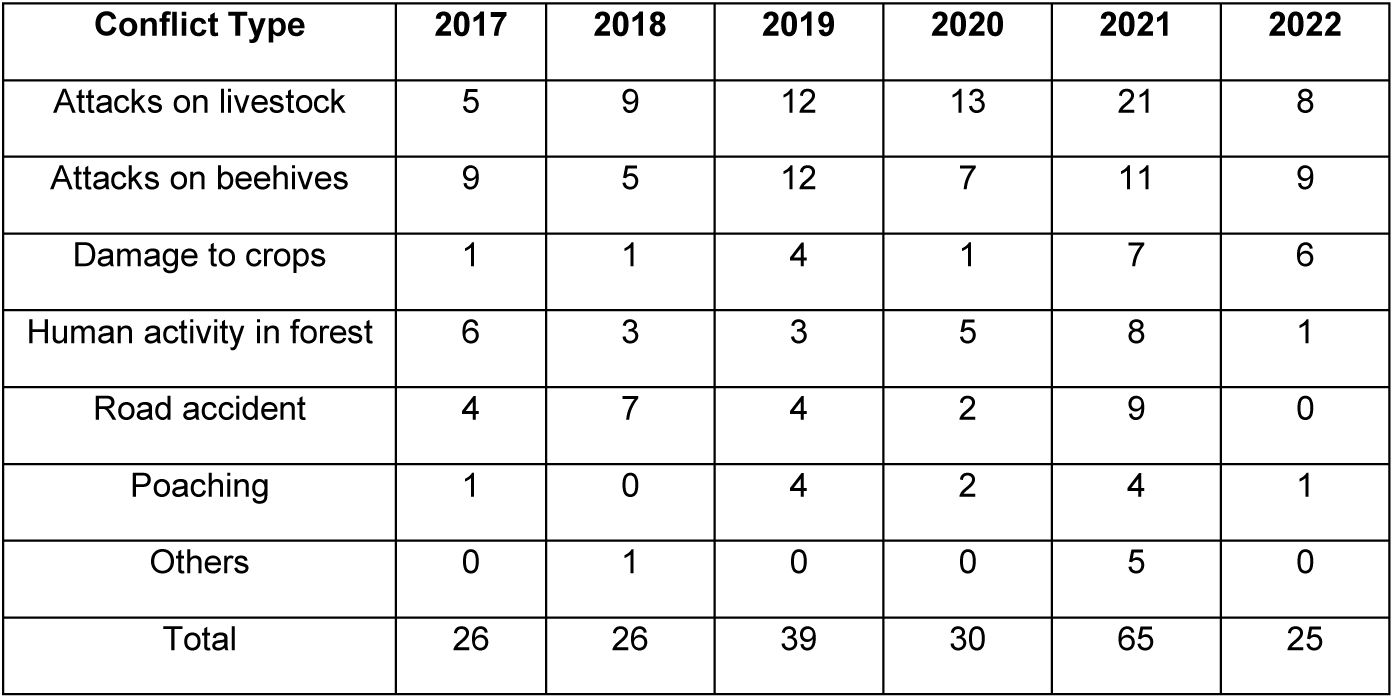
Conflict events’ frequency by the years.

**Table S3:**
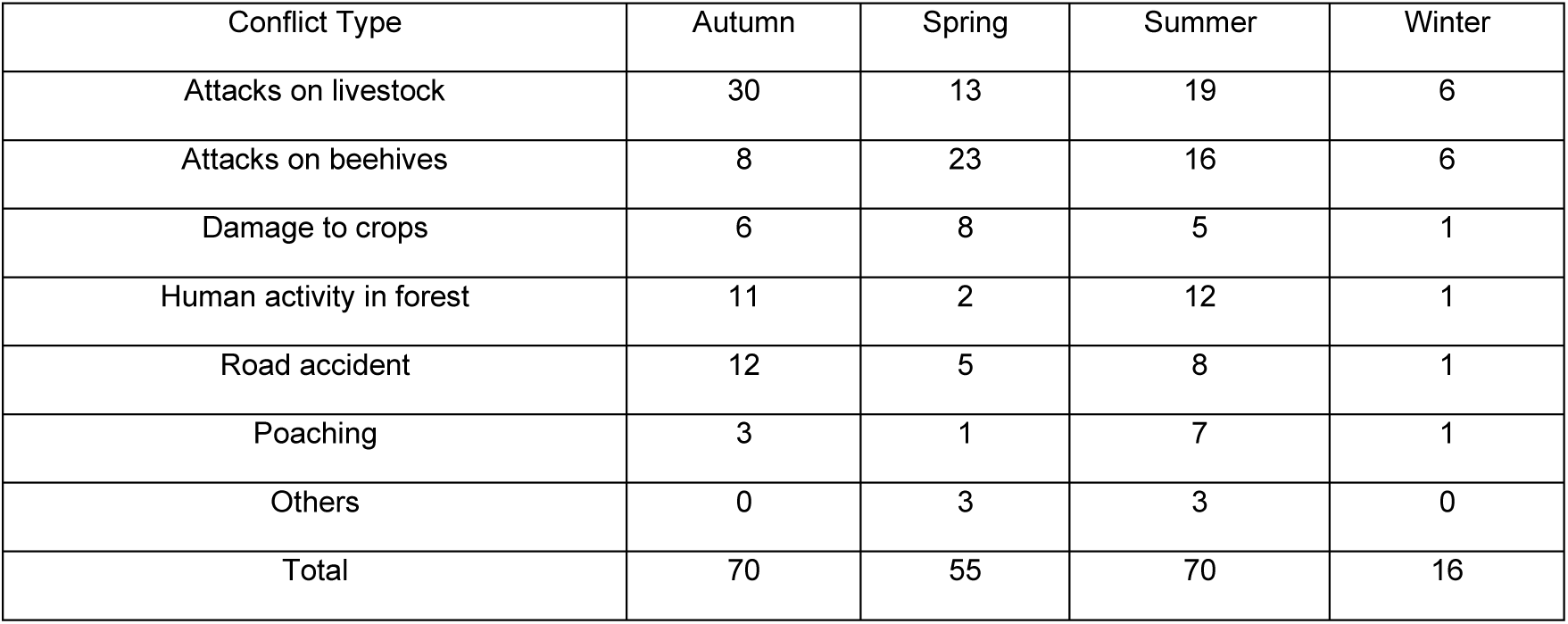
Conflict events’ frequency by the seasons.

**Table S4:**
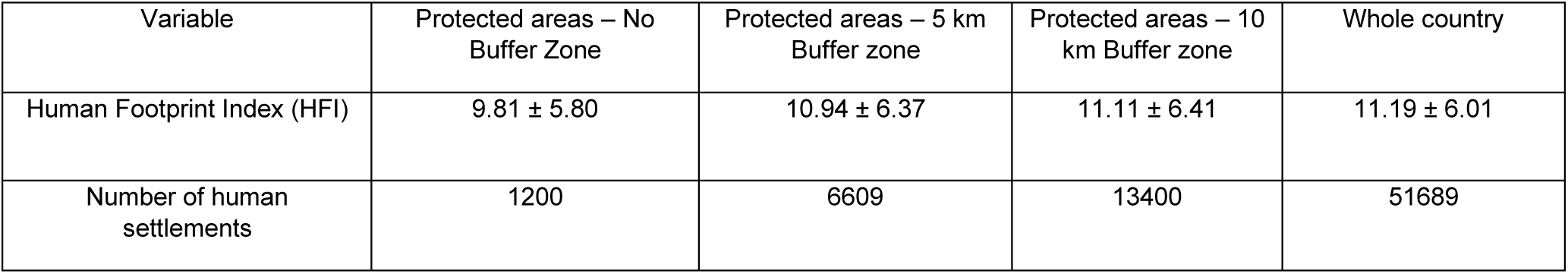
The antropogenic impacts around the protected areas and 5 and 10 km buffer zone around their border.

## Notes

### Competing Interest Statement

The authors have declared no competing interest.

