## Supplementary materials for "Human-brown bear conflicts in Türkiye are driven by increased human presence around protected areas"

**Table S2:** Conflict events' frequency by the years.

**Table S3:** Conflict events' frequency by the seasons.

**Table S4:** The antropogenic impacts around the protected areas and 5 and 10 km buffer zone around their border.

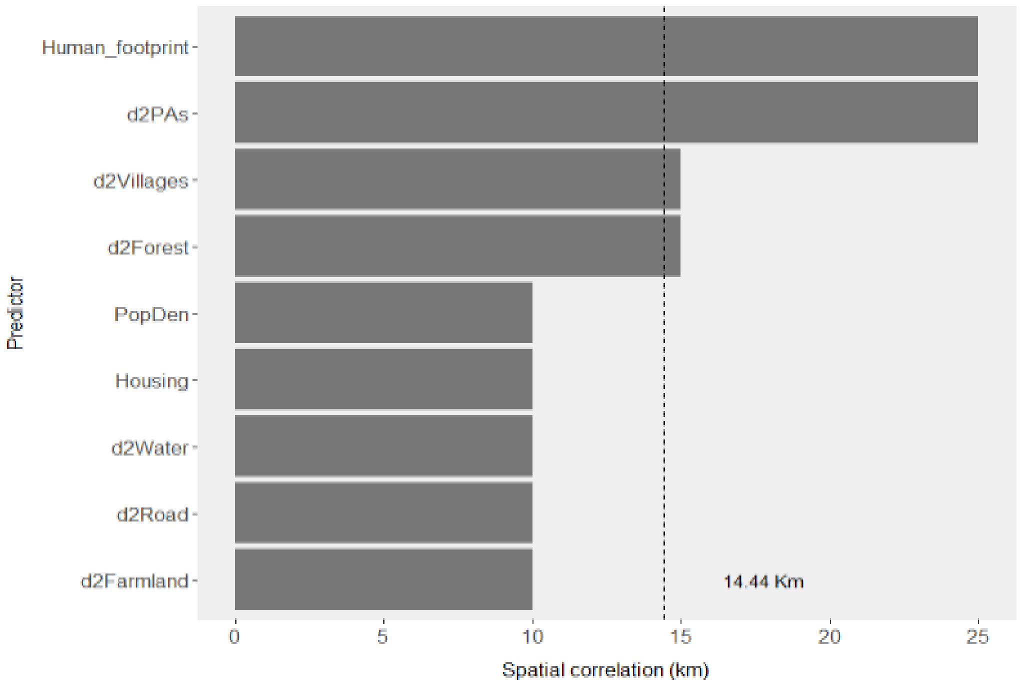


*Figure S1: Minimum and average (dashed line) non-significant autocorrelated distances of variable predictors. d2Farmland: Distance to farmlands; d2Forest: Distance to forests; d2Road: d2PAs: Distance to protected areas; Distance to roads; d2Villages: Distance to villages; d2Water: Distance to waters; Human_footprint: Human Footprint Index; PopDen: Population Density; Housing: Housing Density.*

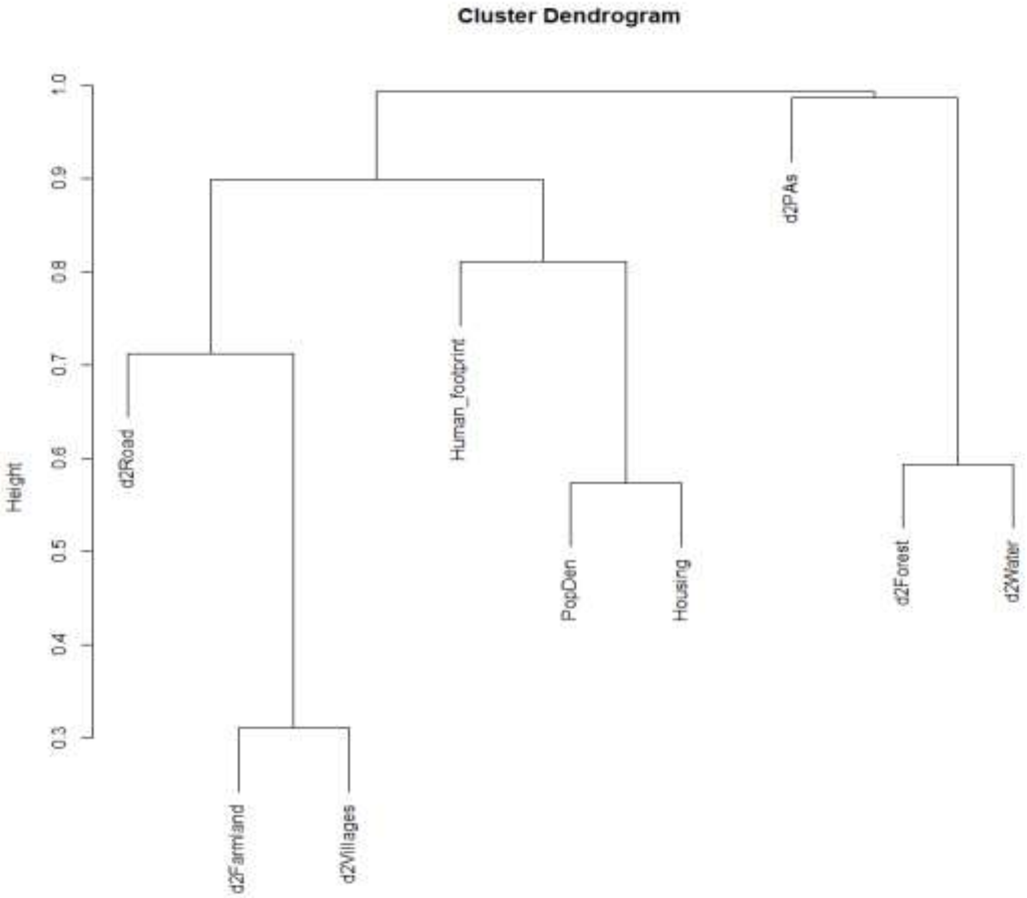


*Figure S2: Cluster dendogram of the environmental variables. d2Farmland: Distance to farmlands; d2Forest: Distance to forests; d2Road: d2PAs: Distance to protected areas; Distance to roads; d2Villages: Distance to villages; d2Water: Distance to waters; Human_footprint: Human Footprint Index; PopDen: Population Density; Housing: Housing Density.*

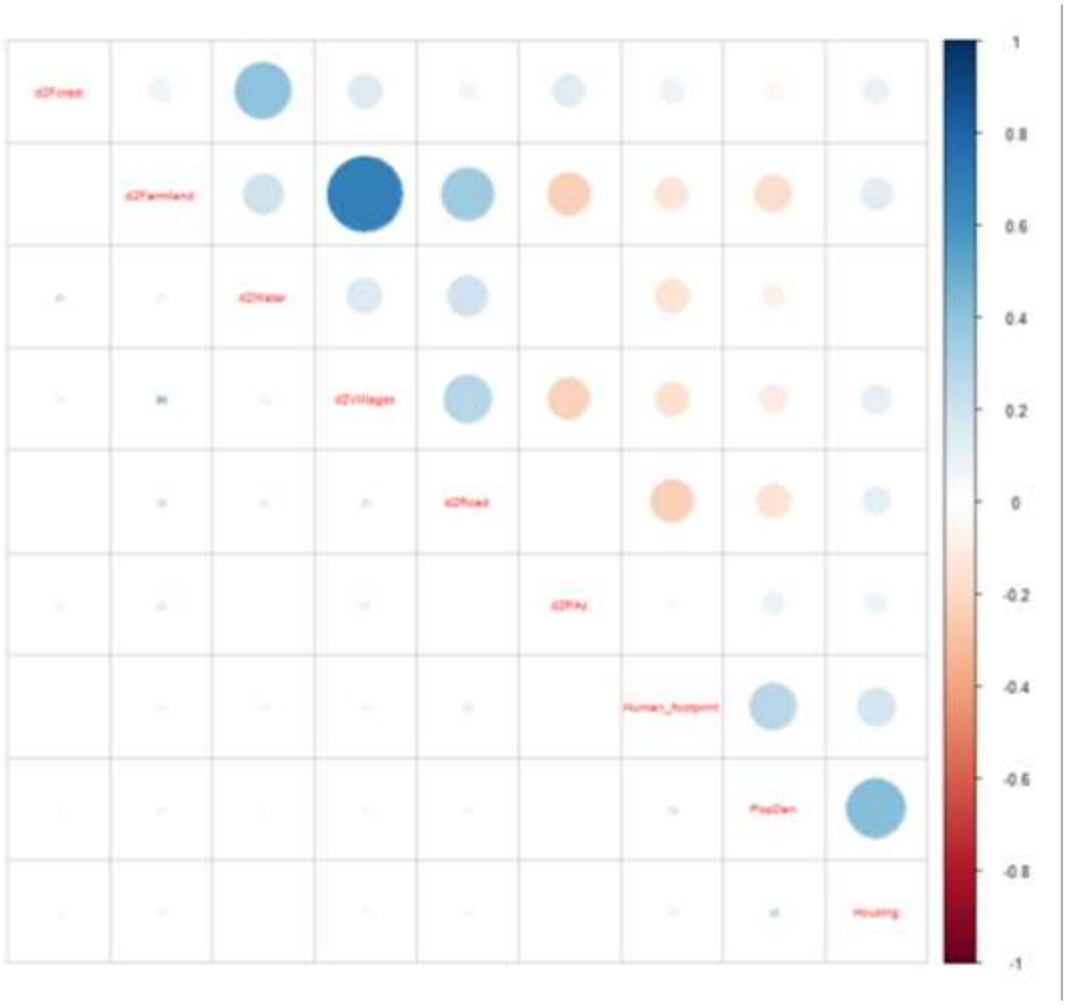


*Figure S3: Correlation matrix of the environmental variables. d2Farmland: Distance to farmlands; d2Forest: Distance to forests; d2Road: d2PAs: Distance to protected areas; Distance to roads; d2Villages: Distance to villages; d2Water: Distance to waters; Human_footprint: Human Footprint Index; PopDen: Population Density; Housing: Housing Density.*

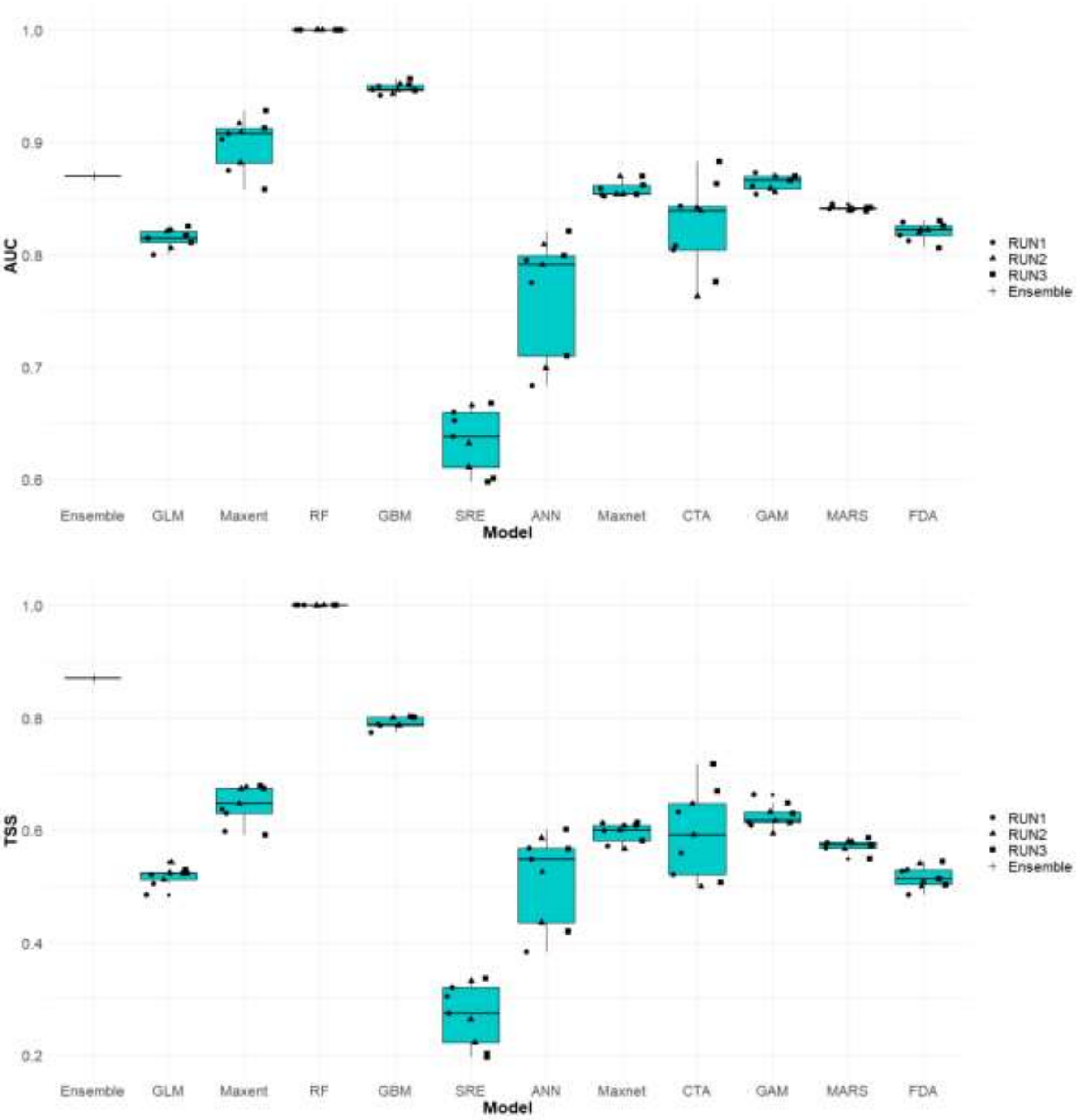


*Figure S4: Receiver operating characteristic (ROC) area under the curve (AUC) (above), and True-skill statistics (TSS) (below) values as predictive performance of different modeling techniques.*

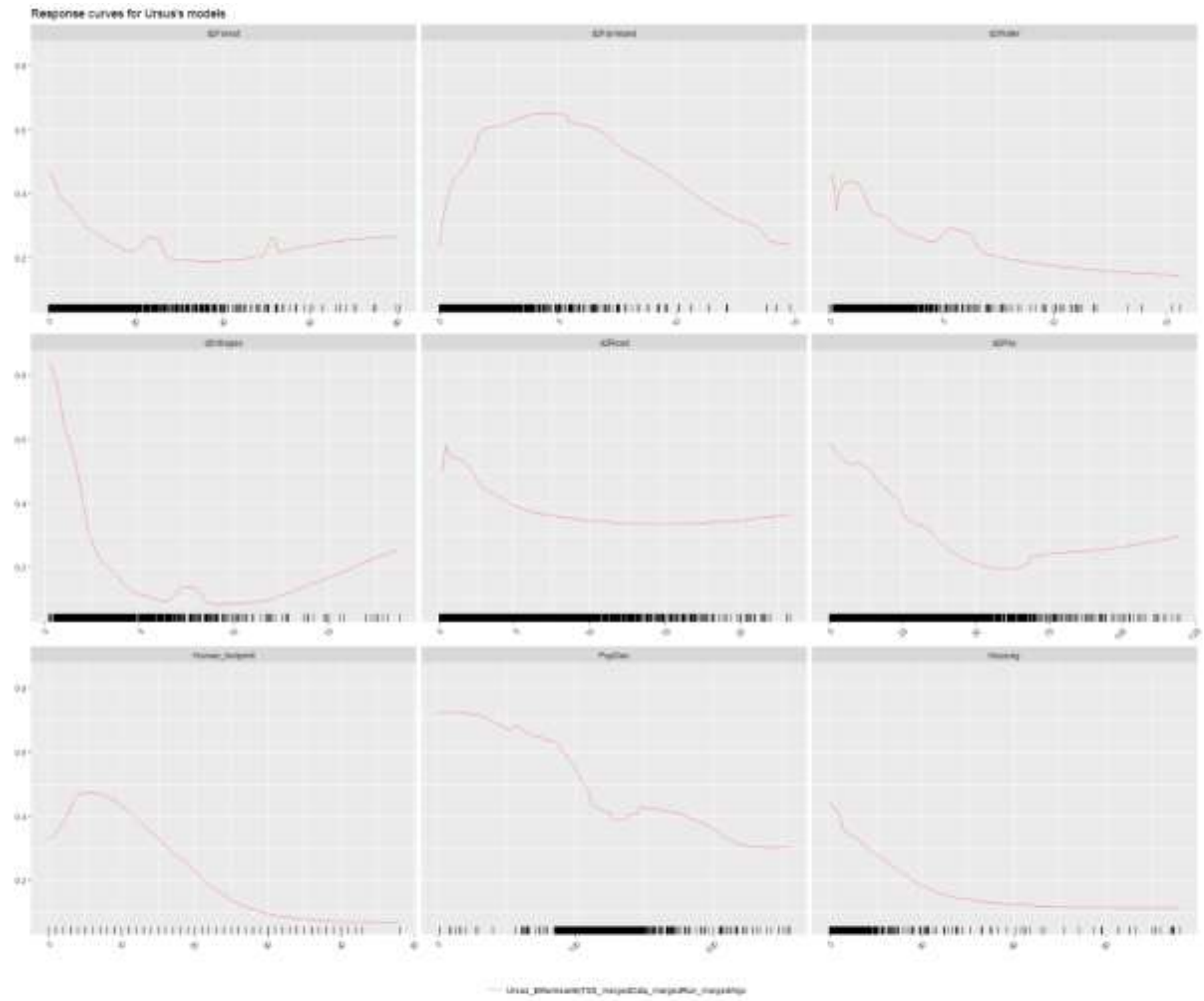


*Figure S5: Response curves of the environmental variables used to risk modeling of human-brown bear conflict across the country. d2Farmland: Distance to farmlands; d2Forest: Distance to forests; d2Road: Distance to roads; d2PAs: Distance to protected areas; d2Villages: Distance to villages; d2Water: Distance to waters; Human_footprint: Human Footprint Index; PopDen: Population Density; Housing: Housing Density.*

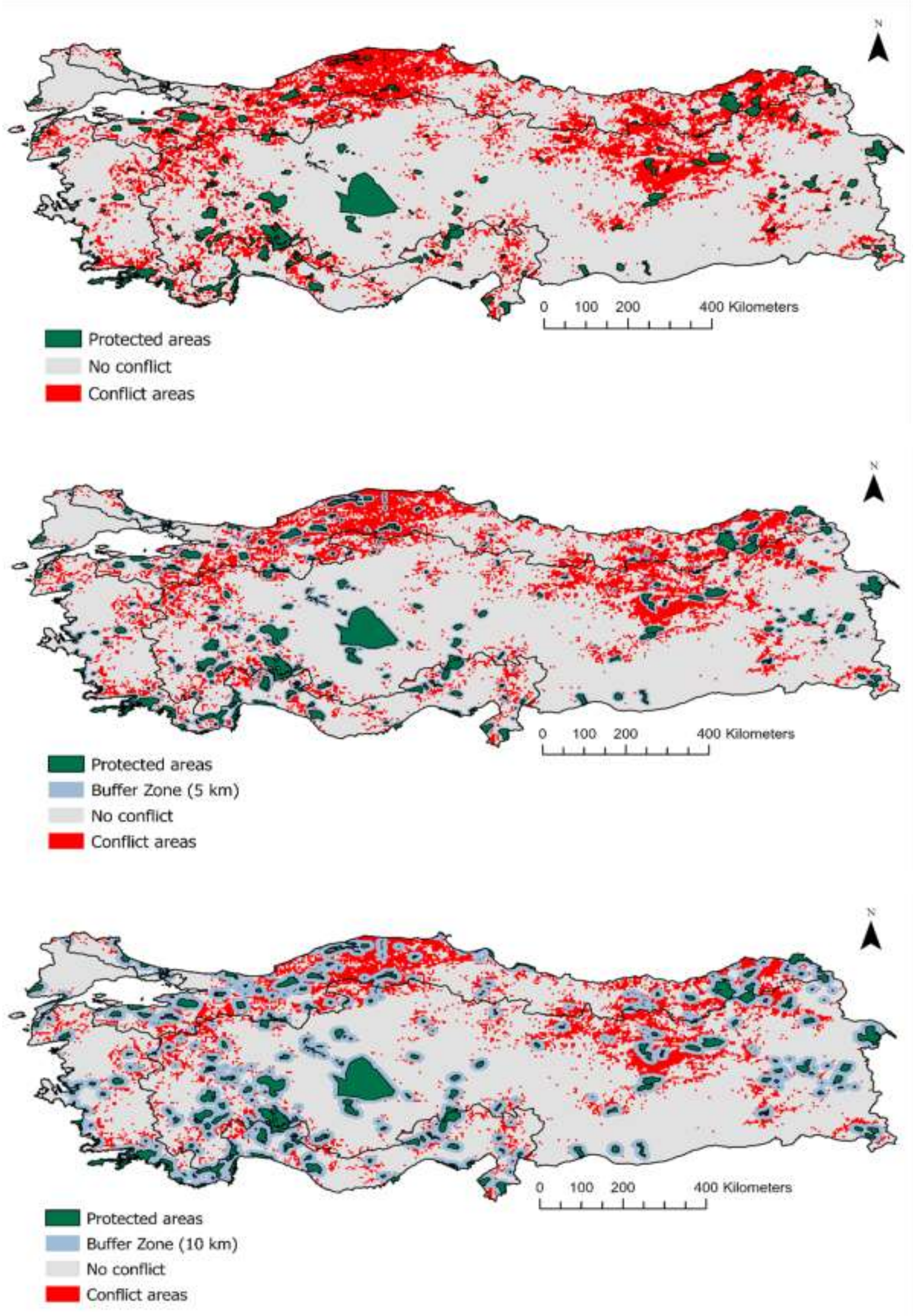


*Figure S6: Binary map of conflict risk across Türkiye based on the ensemble model for 0 (top), 5 (mid) and 10 km buffer zone (bottom) around protected areas.*

*Table S1: Environmental variables used for risk modeling of human-brown bear conflict across Türkiye. All variables were selected to run final model.*

| Source | Variable | Unit | Link |
|---|---|---|---|
| **Corine Land Use Land Cover 2018** | **Distance to Forest** | km | https://land.copernicus.eu/pan-european/corine-land-cover/clc2018 |
| | **Distance to Farmland** | km | |
| **HydroRIVERS – HydroSHEDS** | **Distance to Water** | km | https://www.hydrosheds.org/products/hydrorivers |
| **Open Street Map** | **Distance to road** | Km | https://www.openstreetmap.org/ |
| | **Housing Density** | No. of housing/km$^2$ | |
| **Nature Conservation and Natural Parks** | **Distance to Protected areas** | km | https://www.arcgis.com/apps/View/index.html?appid=5f3978146c4643438ab446620e275269 |
| **SEDAC** | **Human Footprint Index** | - | https://sedac.ciesin.columbia.edu/data/set/wildareas-v3-2009-human-footprint |
| | **Population Density** | No. of persons/km$^2$ | https://sedac.ciesin.columbia.edu/ |
| **-** | **Distance to Villages** | km | - |

*Table S2: Conflict events' frequency by the years.*

| Conflict Type | 2017 | 2018 | 2019 | 2020 | 2021 | 2022 |
|---|---|---|---|---|---|---|
| Attacks on livestock | 5 | 9 | 12 | 13 | 21 | 8 |
| Attacks on beehives | 9 | 5 | 12 | 7 | 11 | 9 |
| Damage to crops | 1 | 1 | 4 | 1 | 7 | 6 |
| Human activity in forest | 6 | 3 | 3 | 5 | 8 | 1 |
| Road accident | 4 | 7 | 4 | 2 | 9 | 0 |
| Poaching | 1 | 0 | 4 | 2 | 4 | 1 |
| Others | 0 | 1 | 0 | 0 | 5 | 0 |
| Total | 26 | 26 | 39 | 30 | 65 | 25 |

*Table S3: Conflict events' frequency by the seasons.*

| Conflict Type | Autumn | Spring | Summer | Winter |
|---|---|---|---|---|
| Attacks on livestock | 30 | 13 | 19 | 6 |
| Attacks on beehives | 8 | 23 | 16 | 6 |
| Damage to crops | 6 | 8 | 5 | 1 |
| Human activity in forest | 11 | 2 | 12 | 1 |
| Road accident | 12 | 5 | 8 | 1 |
| Poaching | 3 | 1 | 7 | 1 |
| Others | 0 | 3 | 3 | 0 |
| Total | 70 | 55 | 70 | 16 |

*Table S4: The antropogenic impacts around the protected areas and 5 and 10 km buffer zone around their border.*

| Variable | Protected areas – No Buffer Zone | Protected areas – 5 km Buffer zone | Protected areas – 10 km Buffer zone | Whole country |
|---|---|---|---|---|
| Human Footprint Index (HFI) | 9.81 ± 5.80 | 10.94 ± 6.37 | 11.11 ± 6.41 | 11.19 ± 6.01 |
| Number of human settlements | 1200 | 6609 | 13400 | 51689 |
